# Neural Mechanism of Hunger-gated Food-seeking and Evaluating

**DOI:** 10.1101/2020.10.23.352187

**Authors:** Young Hee Lee, You Bin Kim, Kyu Sik Kim, Ha Young Song, Mirae Jang, Dong-Soo Ha, Joon Seok Park, Sang-Ho Jung, Jaegeon Lee, Kyung Min Kim, Deok-Hyeon Cheon, Inhyeok Baek, Min-Gi Shin, Eun Jeong Lee, Sang Jeong Kim, Hyung Jin Choi

**Author notes:** These authors contributed equally (Young Hee Lee, You Bin Kim, Kyu Sik Kim, Ha Young Song, Mirae Jang). Correspondence and requests for materials should be addressed to (H.J.C.) and (S.J.K.).

## Abstract

The physiological need for energy evokes motivated feeding behaviours that help to ensure survival. However, the neural mechanisms underlying the generation of food motivation remain poorly understood. We investigated these mechanisms by subdividing feeding-related motivated behaviours into food-seeking, evaluating, and swallowing. Micro-endoscopic results indicated that neurons containing leptin receptors (LepRs) in the lateral hypothalamus (LH) are the major food-specific subpopulation of LH^GABA^ neurons. Optogenetic manipulation of LH^LepR^ neurons bidirectionally regulated both food-seeking and evaluating. Furthermore, micro-endoscope data revealed that distinct LH^LepR^ neurons encode seeking and evaluating. Computational modelling analysis demonstrated that LH^LepR^ neurons encode motivation, whereas neurons containing agouti-related peptide and neuropeptide Y (AgRP/NPY) encode the need for food. Additionally, slice studies revealed that NPY decreases inhibitory input to LH^LepR^ neurons via LH^GABA^ interneurons. This mechanism explains the permissive gate role of hunger (food need) in seeking/evaluating motivation. Together, the present study provides a comprehensive neural mechanism of how physiological needs drive distinct motivated behaviours.

## Introduction

To maintain homeostasis, physiological need drives the diverse motivations required to evoke appropriate behaviours^1,2,3^. In the context of feeding, motivated behaviours are divided into “seeking” (moving toward the target), “evaluating” (examining the target by tasting, smelling, etc.), and “swallowing” (consuming the target)^45,6,7^. These separate food-motivated behaviours are orchestrated by exteroceptive and interoceptive signals in the brain to execute the appropriate feeding behaviours required for survival^4, 8^. To study the precise mechanisms orchestrating these distinct motivations, we developed specific behavioural experimental paradigms to induce and distinguish the three feeding behaviours.

Heterogenous GABAergic neurons are widely distributed in the lateral hypothalamus (LH) and are involved in various motivated behaviours, including feeding^9,10,11,12,13^. Among LH^GABA^ neurons, leptin receptor (LepR) neurons are reported to be associated with feeding behaviour, although the results of previous studies remain controversial^14,15,16,17,18^. To investigate whether distinct neural populations separately encode motivation for food-seeking and/or evaluating, we focused on LH^GABA^ and LH^LepR^ neurons using these newly developed behavioural paradigms. Next, we investigated how neurons containing agouti-related peptide and neuropeptide Y (AgRP/NPY), which are well known to regulate feeding, orchestrate motivation for food-seeking/evaluating^19,20,21^. Neural dynamics and causal perturbations observed in these experiments revealed that two distinct LH^LepR^ populations separately motivate seeking and evaluating during the state of hunger, and this process is gated by AgRP/NPY neurons that encode the need for food.

## Results

### LH^LepR^ neurons are food-specific subpopulation of LH^GABA^ neurons

To investigate food-specific neurons within the population of LH^GABA^ neurons, three food context tests and one non-food context test were performed using a micro-endoscope (Fig. 1a, c, e). The results for 218 LH^GABA^ neurons showed that only a minor subpopulation of neurons (8%) was food specific (Fig. 1e, g-j). These findings imply the existence of an isolated food-specific population within the vast population of LH^GABA^ neurons. Since leptin decreases food motivation by inhibiting LH^LepR^ neurons^14, 15, 18^, and LH^LepR^ neurons have been reported to be associated with feeding^16, 18^. Therefore, we hypothesised that LH^LepR^ neurons may represent this food-specific subpopulation.

**Fig. 1.**
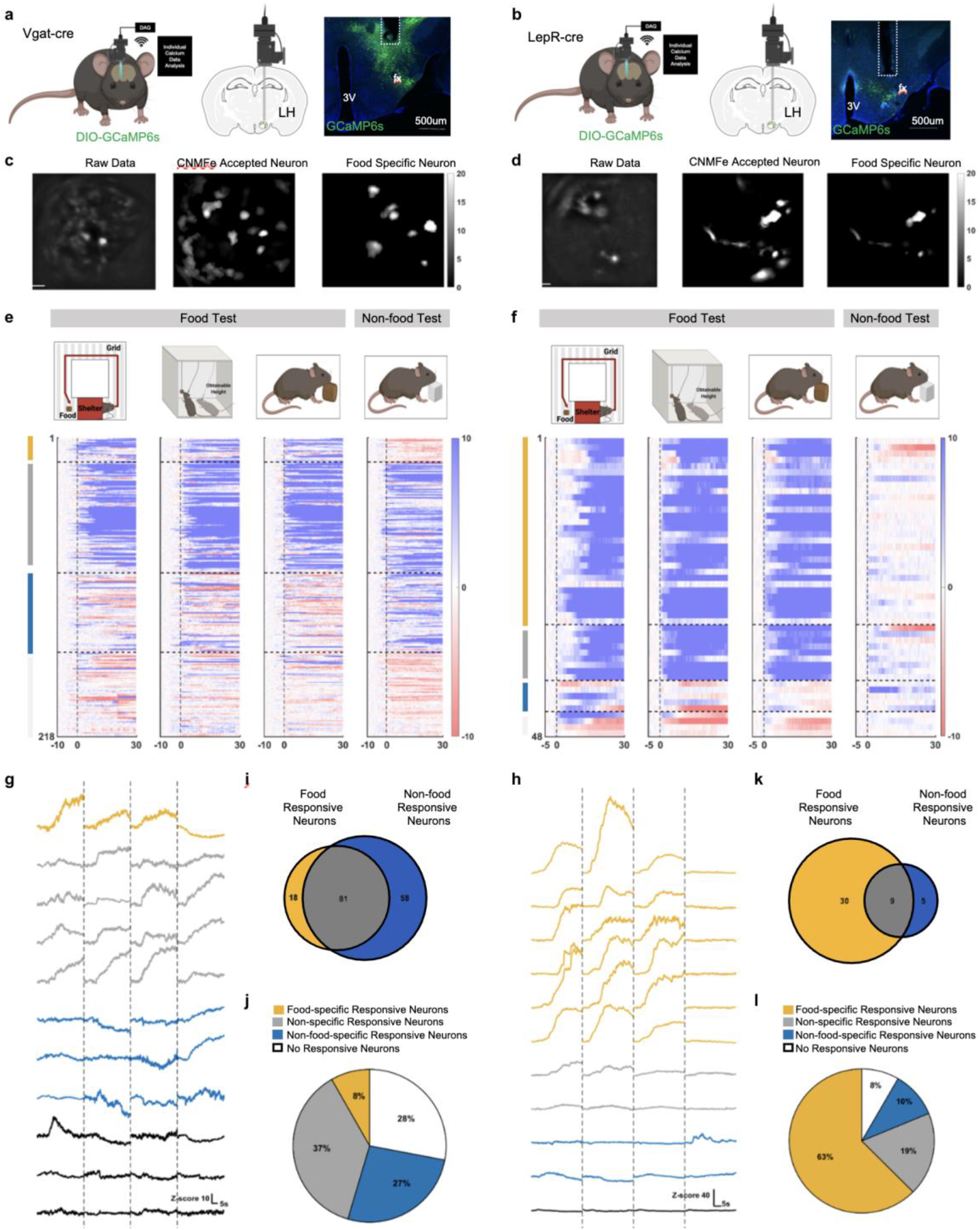
LH^LepR^ neurons are the food-specific subpopulation of LH^GABA^ neurons. **a, b,** Schematic of micro-endoscopic calcium imaging (left, middle), and image of GCaMP6s expression in the LH from Vgat-Cre and LepR-Cre mice (right). **c, d,** Spatial map of raw data (left), accepted cells using CNMFe (middle), and cells that only respond to food-related behaviour (right), from Vgat-cre (**c**) and LepR-cre (**d**) mice. Cells are coloured according to the maximum Z-score. **e, f,** Schematic of the voluntary behaviour maze test, vertically placed food test, food test, and non-food test (top). Heatmap depicting calcium signals that respond to each test (below). Four populations are discriminated: food-specific responsive (yellow), non-specific responsive (gray), non-food-specific responsive (blue), and non-responsive (white) cells. (LH^GABA^ neurons 218 cells, 5 mice, LH^LepR^ neurons 48 cells, 4 mice). **g, h,** Representative traces of four populations. **i, k,** Venn diagram of food-specific responsive. LH^GABA^ neurons 8% (18/ 218 cells) (**i**), LH^LepR^ neurons 63% (30/total 48 cells) (>4σ) (**k**). **j, l,** Proportion of food-specific responsive (yellow), non-specific responsive (gray), non-food-specific responsive (blue), and non-responsive (white) cells.

To identify the distribution of LepR neurons in the LH, we performed whole-LH three-dimensional (3D) tissue clearing (Supplementary Video 1) and 2D histological mapping using LepR-tdTomato mice (Extended Data Fig. 1a–k). Our results indicated that the LH^LepR^ neurons were mainly distributed in the middle (−1.5 mm from bregma) and posterior (−2.2 mm from bregma) regions. Based on these distribution data, we targeted LepR neurons in the middle region of the LH. LH^LepR^ neural activity was measured using a micro-endoscope (Fig. 1b, d, f). Most LH^LepR^ neurons (63%) were specific to the food context (Fig. 1h, k,l), and most LH^LepR^ neurons did not respond to water (Extended Data Fig. 2a–e).

Based on the available single-cell RNA sequencing data for the LH^22^, we discovered that LH^LepR^ neurons comprise only 4% of LH^GABA^ neurons (Extended Data Fig. 1m). A previous study has also reported that LH^LepR^ neurons comprise <20% of LH^GABA^ neurons^15^. Although LH^LepR^ neurons represented only a minor portion (4–20%) of LH^GABA^ neurons, most LH^LepR^ neurons were food-specific (63%), in contrast to findings for LH^GABA^ neurons (8%). Collectively, these results suggest that LH^LepR^ neurons are the major food-specific neurons among LH^GABA^ neurons (Extended Data Fig. 2k).

### LH^LepR^ neurons are activated during food-seeking and evaluating behaviors

To identify the temporal dynamics of LH^LepR^ neurons in food-seeking and evaluating behaviours, we performed various food-related tests using fibre photometry (Fig. 2a-b). LH^LepR^ neural activity significantly increased at each feeding bout with specific time-locked temporal dynamics (Fig. 2c-g, Extended Data Fig. 2f–j). Interestingly, LH^LepR^ neural activity increased even before physical contact with food, implying that LH^LepR^ neurons may be involved in seeking or evaluating. To provide sufficient temporal distinction between seeking and evaluating, we designed tests specific to each behaviour (Fig. 2h, k). Before conditioning, since mice were not aware of the food location, they explored the whole maze (non-goal-directed locomotion) (Fig. 2i). LH^LepR^ neural activity did not increase during this non-goal-directed locomotion (Fig. 2l). LH^LepR^ neural activity started to increase when mice evaluated food at the end of the corridor. However, after conditioning, the mice moved directly to the food at the end of the corridor (goal-directed seeking; significantly shorter time to food contact) (Fig. 2j). When compared with that observed before conditioning, LH^LepR^ neural activity started to increase significantly when mice initiated food-seeking, and this increase was sustained during the evaluating stage (Fig. 2m). Additional tests revealed that LH^LepR^ neural activity decreased when mice voluntarily terminated both seeking and evaluating, suggesting that LH^LepR^ neural activity is significantly correlated with these behaviours (Extended Data Fig. 3a-j).

**Fig. 2.**
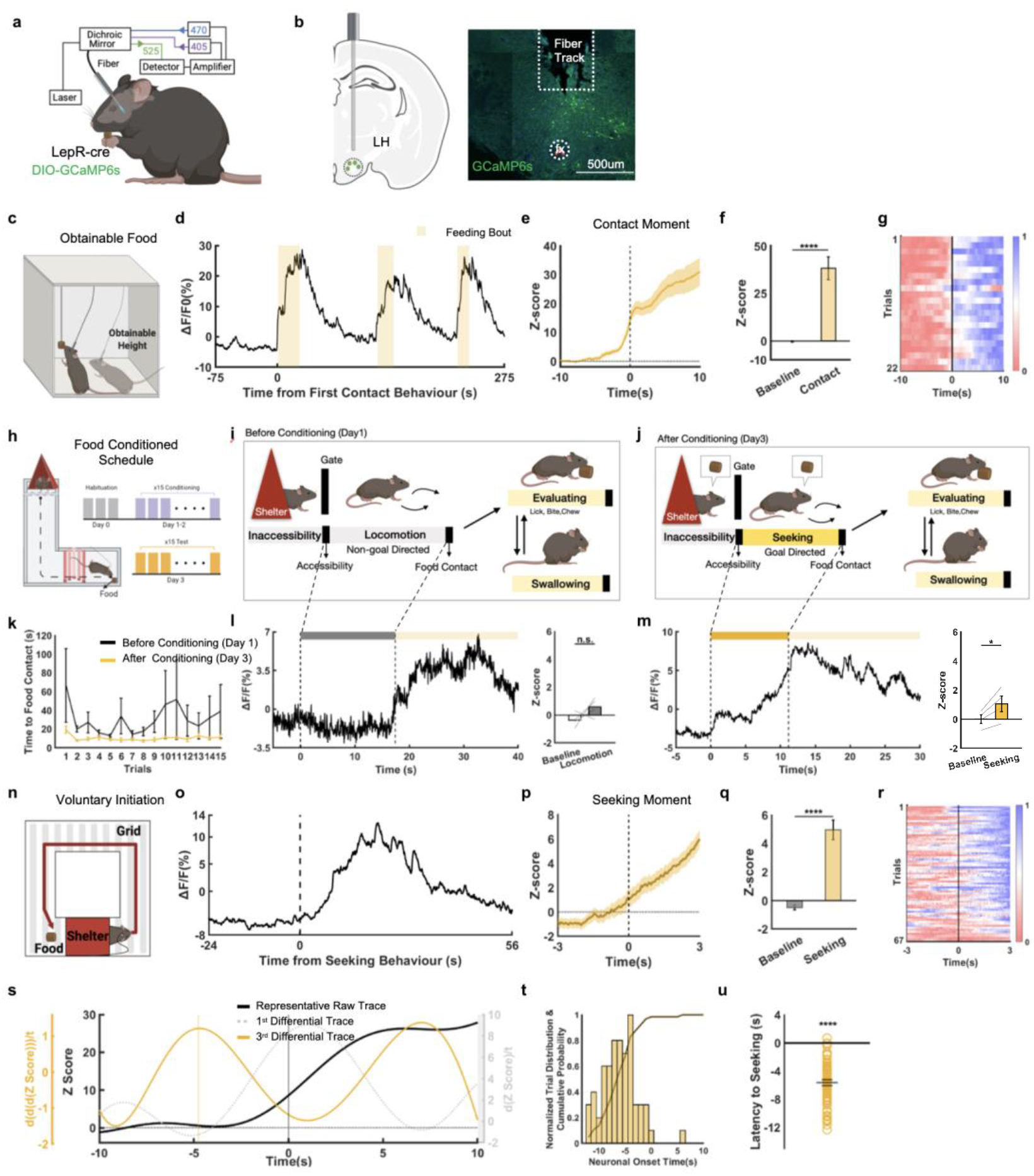
Activity of LH^LepR^ neurons is time-locked to food-seeking and evaluating Behaviours. **a,** Schematic of virus injection/fibre insertion for fibre photometry in the LH from LepR-Cre mice. **b,** A representative image validates ChR2 expression in LepR neurons and optical fiber tract above the LH. **c**, Schematic of the vertically placed food test (obtainable). **d, e,** Representative calcium traces **(d)** and average of Z-score from LH^LepR^ calcium response **(e)** aligned to contact the food. Yellow shaded box: from the moment of contact behaviour to end of food consumption. (5 mice, 22 trials). **f,** Quantification of Z-score in calcium signal change from **(e)**. Comparison between the baseline (−8 to −7s) and after contact (9 to 10s). **g**, Heatmap depicting normalized LH^LepR^ neural activity aligned to contact moment of the food. **h,** Schematic of the food conditioned maze test. **i, j,** Phases of dynamic feeding before conditioning (**i**) and after conditioning (**j**). **k**, Time from food accessibility to food contact, before and after conditioning. **l, m,** Representative calcium signal of LH^LepR^ neurons aligned to food accessibility (left) and quantification of Z-score in calcium signal change (right) before (**l**) and after (**m**) conditioning. **n**, Schematic of the voluntary behavioural maze test. **o, p,** Representative of calcium traces (**o**) and average Z-score (**p**) from LH^LepR^ neural calcium signal response aligned to voluntary seeking initiation moment. (4 mice, 67 trials). **q**, Quantification of Z-score in calcium signal change from (**p**). Comparison between baseline (−8 to −7 s) and after search onset (2 to 3 s). **r**, Heatmap depicting normalised LH^LepR^ calcium signal aligned to voluntary seeking behaviour. **s**, Representative neural onset of LH^LepR^ calcium signal during (**n**). **t, u**, Cumulative probability distribution (**t**) and histogram (**u**) of LH^LepR^ neural onset during (**n**) (4 mice, 67 trials). Neural onset occurred at −5.623 ± 0.418 s. Data are mean ± s.e.m. See Supplementary Table 1 for statistics.

To precisely quantify the temporal onset of LH^LepR^ neural activity and compare it with the voluntary onset of food-seeking, we allowed mice to make a voluntary decision regarding when to initiate food-seeking behaviour. The mice were trained in the doorless shelter with an appropriate amount of electrical shock and food reward, such that they intermittently made a voluntary decision to move out of the shelter to seek the food at the end of the corridor (Fig. 2n). LH^LepR^ neural activity began to increase significantly before the mice voluntarily initiated seeking (Fig. 2o-r, Supplementary Video 2). Furthermore, the onset of LH^LepR^ neural activity significantly preceded the onset of seeking by an average of approximately 6 s (Fig. 2s-u). These results indicate that LH^LepR^ neurons exhibit a causal temporal relationship with food-seeking behaviour, suggesting that activity in LH^LepR^ neurons represents the motivation for seeking, not the consequence of seeking behaviour.

### Activation of LH^LepR^ neurons evoke food-seeking and evaluating

To investigate whether the activation of LH^LepR^ neurons increases food-seeking or evaluating behaviour (Fig. 3a, b), we displayed two foods and two non-food objects with similar appearance in the arena in an alternative order and measured behavioural changes during LH^LepR^ neural activation (Extended Data Fig. 4a). Activation of LH^LepR^ neurons significantly increased the distance moved, velocity in the seeking zone, the frequency of visiting the evaluating zones, and the time spent in the evaluating zones (Extended Data Fig. 4b-g).

**Fig. 3.**
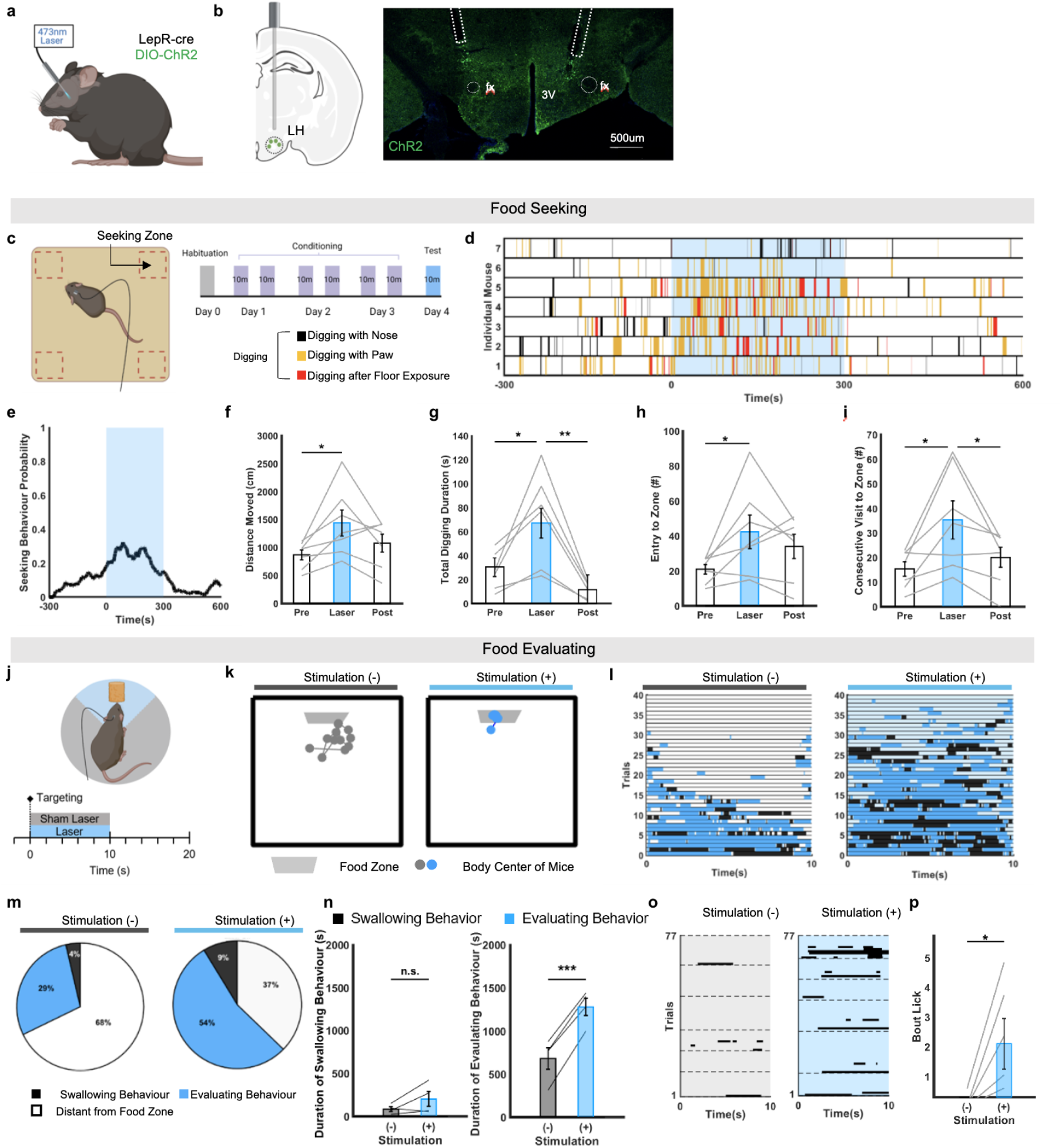
Activation of LH^LepR^ neurons drives motivation for food-seeking and evaluating. **a, b**, Schematic of optogenetic activation and image of ChR2 expression in LH^LepR^ neurons. **c**, Schematic of the hidden food-seeking test. **d**, Raster plot during (**c)** (7 mice). **e**, Behavioural probability from (**d**). **f-i**, Quantification of distance moved (**f**), total digging duration (**g**), frequency of food zone entry (**h**) and consecutive visit to food zone (**i**). **j**, Schematic of the nearby food-evaluating test. **k**, Trajectory of centroid of snout, left hand, and right hand in bottom view. **l,** Raster plot of during (**j**). **m**, Proportion of swallowing and evaluating from (**l**) (4 mice). **n**, Quantification of swallowing and evaluating from (**l**) (4 mice). **o**, Raster plot of evaluating bout duration (6 mice). **p**, Quantification of evaluating (6 mice).

To scrutinise the role of LH^LepR^ neurons in the regulation of food-seeking, mice were conditioned to food hidden in the four corners of an open-field chamber filled with bedding (Fig. 3c, Supplementary Video 3). On the photostimulation day, the mice were placed in the same area covered with bedding without food to evoke seeking behaviour only, without any evaluating. Activation of LH^LepR^ neurons significantly increased seeking behaviours (digging with nose, digging with paw, and digging after floor exposure), entry into the seeking zones, and locomotion (Fig. 3d-i). These results indicate that activation of LH^LepR^ neurons evokes seeking behaviour. Consistent with the activation of middle LH^LepR^ neurons, chemogenetic activation of posterior LH^LepR^ neurons also significantly evoked seeking behaviours (Extended Data Fig. 4p-r).

To precisely investigate evaluating behaviour and distinguish it from swallowing, the mice were confined to a small chamber to minimise seeking (Fig. 3j). To accurately distinguish evaluating behavior from swallowing behavior, we performed two independent tests using different behavior analysis methods. First, we used a deep-learning based animal pose estimation method (DeepLabCut)^23^. The results of this analysis revealed that stimulating LH^LepR^ neurons significantly evoked only evaluating behaviors and not swallowing behavior (Fig. 3k-n, Supplementary Video 4). Second, by using a manual behavior analysis method, we revealed that stimulating LH^LepR^ neurons significantly evoked evaluating behaviors (Fig. 3o, p, Extended Data Fig.4h-l). These results demonstrate that the stimulation of LH^LepR^ neurons evokes only food-evaluating but not swallowing behavior.

Additionally, to investigate the valence effect of LH^LepR^ neurons, real-time place preference was performed (Extended Data Fig. 4m). The results showed that mice significantly changed their preference to the laser stimulation side (Extended Data Fig. 4n middle), in accordance with previous results^16, 24^. When the laser stimulation side was reversed in the last session, the preference tends to change according to the stimulation side (Extended Data Fig. 4n right, 4o). These results indicate that the activation of LH^LepR^ neurons has a positive valence effect.

### Inhibition of LH^LepR^ neurons attenuates food-evaluating behavior

To examine whether the inhibition of LH^LepR^ neurons decreases evaluating behaviours, halorhodopsin was expressed in LH^LepR^ neurons (Fig. 4a). To quantify the food evaluating behavior of mice, we used a sequential behavioural paradigm to examine both evaluating and swallowing behaviours when multiple small snacks were presented during several interleaved photoinhibition blocks (Fig. 4b, Supplementary Video 3). During the sessions, mice sequentially evaluated the food (savoring it by bringing one piece of the snack to the mouth), after which they swallowed it (Fig. 4c). Inhibition of LH^LepR^ neurons significantly decreased evaluating behaviour, after adjusting for the satiation factor (Fig. 4d). Inhibition of LH^LepR^ neurons did not significantly alter the frequency at which evaluating behaviours were initiated (Fig. 4e). However, inhibition of LH^LepR^ neurons significantly decreased the duration of each evaluating bout, number of completions (completing snack swallowing), and the number of long evaluating bouts (>3s) (Fig. 4f-n). These findings indicate that photoinhibition of LH^LepR^ neurons disturbs the maintenance of evaluating behaviour and results in premature termination of such behaviour.

**Fig. 4.**
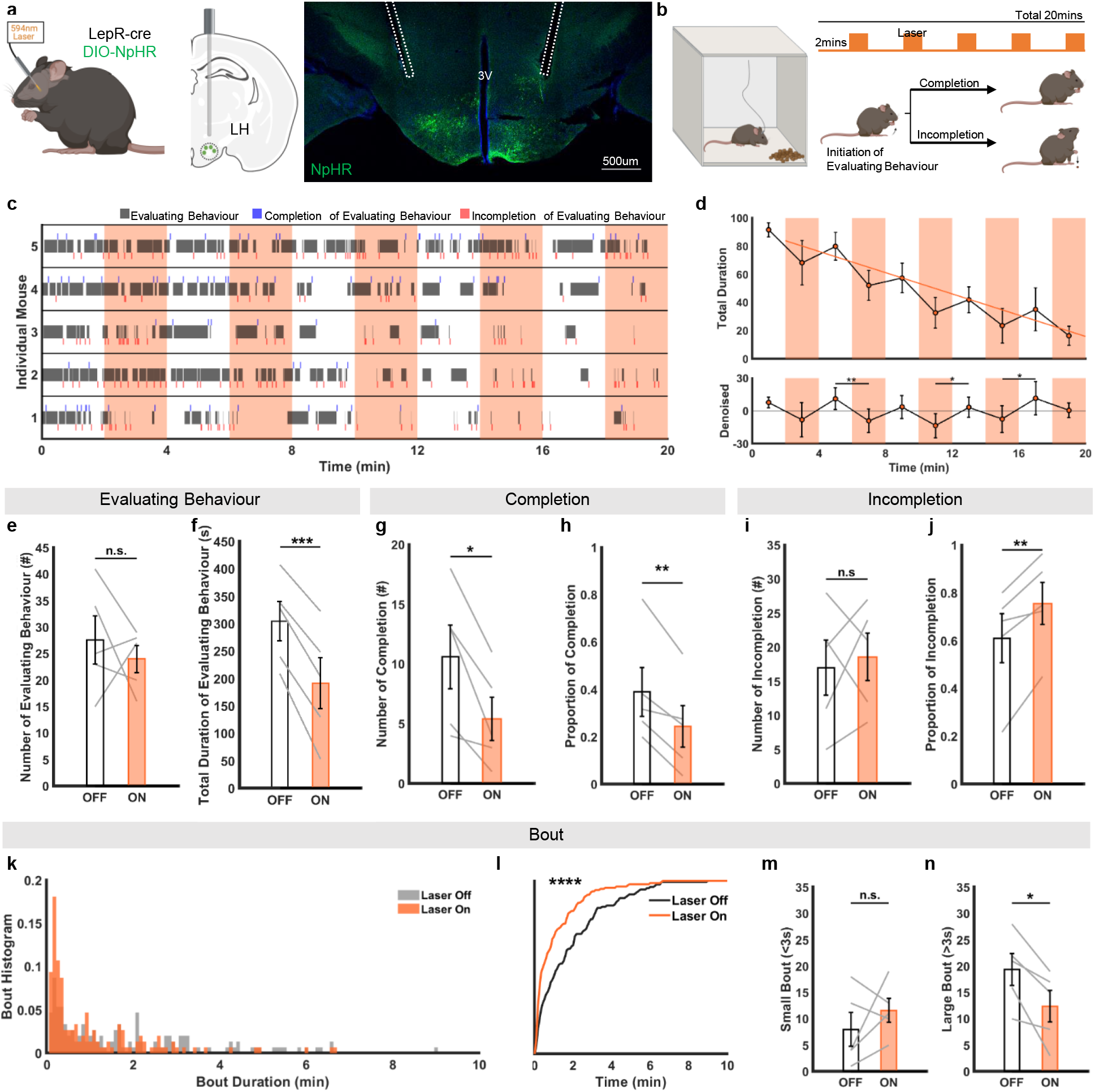
Inhibition of LH^LepR^ neurons decreases motivation for food-evaluating. **a,** Schematic of optogenetic inhibition (left) and image of NpHR expression in LepR neurons in LH (right). **b,** Schematic of the multiple food-evaluating test. **c,** Raster plot of evaluating during (**b**) (5mice). **d**, Average duration of evaluating behaviour (top). Calibrated graph (bottom) of top. **e-h**, Quantification from (**c**) frequency (**e**), total duration of evaluating (**f**), frequency (**g**), proportion of completion of evaluating (**h**). **i-j,** frequency (**i**), and proportion of incompletion evaluating (**j)**. **k, l**, Histogram (**k**) and cumulative probability distribution (**l**) of evaluating behaviour bout from (**k**). **m, n**, Quantification of frequency of small bout (<3s), (**m**) and large bout (>3s), (**n**). Data are mean ± s.e.m. See Supplementary Table 1 for statistics.

### Two distinct subpopulations of LH^LepR^ neurons individually encode food-seeking and evaluating behaviors

To evaluate heterogeneity among populations of LH^LepR^ neurons, we separately investigated changes in LH^LepR^ neural activity during food-seeking and evaluating using a micro-endoscope (Fig. 5a). To distinguish between seeking and evaluating, we modified the voluntary behavioural maze test described above (Fig. 2n). During food sessions, mice sequentially performed seeking and evaluating (Fig. 5b left). In contrast, during no-food sessions, mice performed seeking without evaluating (Fig. 5b right, Supplementary Video 5). We identified two distinct neural populations that specifically and consistently responded to seeking or evaluating (Fig. 5c). Activity in one population of neurons only increased during seeking and not during evaluating (seeking neurons) (Fig. 5f-i, Extended Data Fig. 5a). Another population of neurons only increased during evaluating (evaluating neurons) (Fig. 5j-m, Extended Data Fig. 5b). The two populations were distinctively separated in a 3D scored plot (Fig. 5d). Among the population of LH^LepR^ neurons, 25% were seeking neurons, and 39% were evaluating neurons (Fig. 5e). These results indicate that two distinct LH^LepR^ neural populations separately encode seeking and evaluating.

**Fig. 5.**
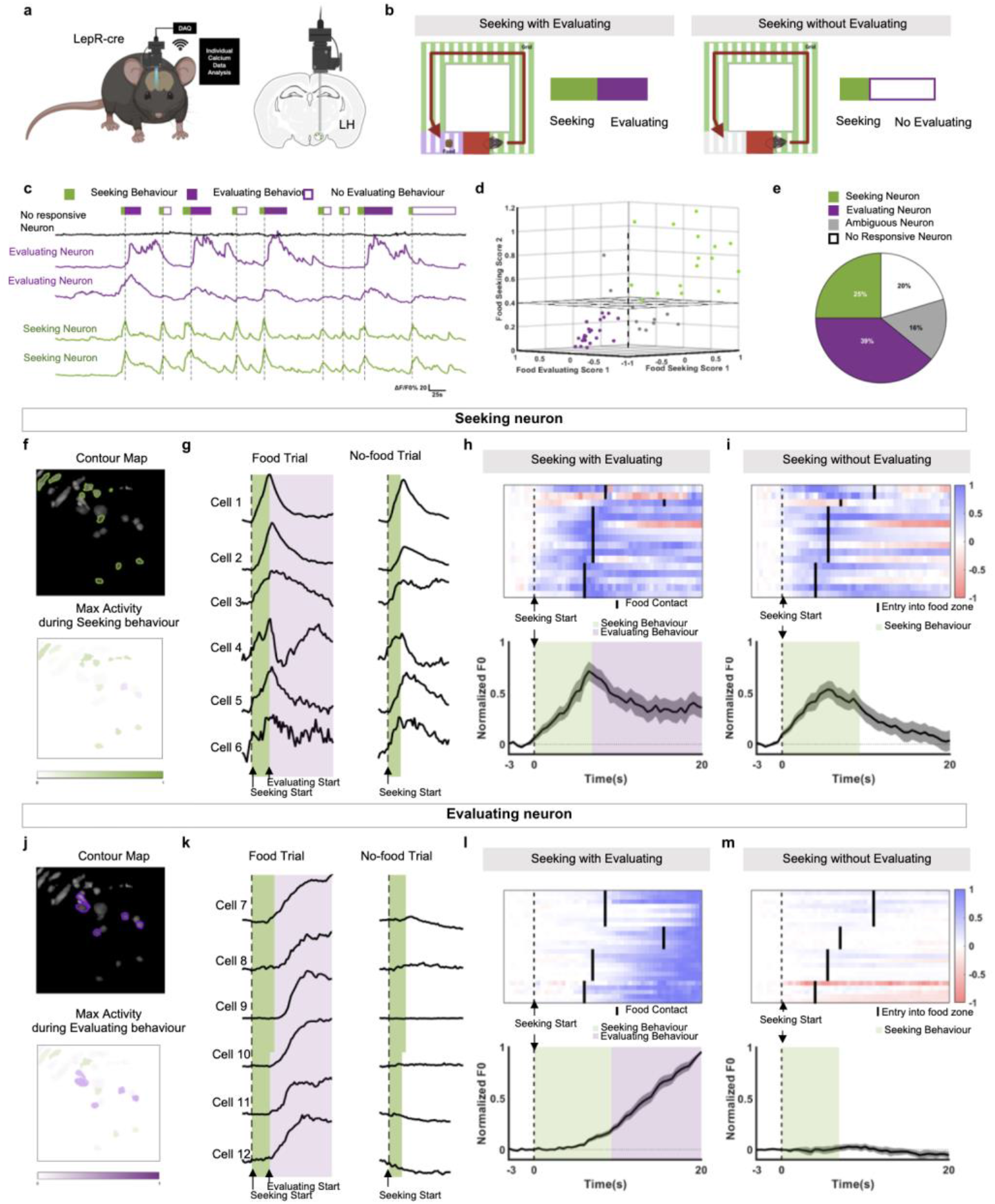
Two distinct populations of LH^LepR^ neurons encode food-seeking and evaluating motivation. **a**, Schematic of virus injection/GRIN lens insertion for micro-endoscopic calcium imaging in the LH from LepR-Cre mice. **b**, Schematic of the voluntary behavioural maze test. Seeking with evaluating condition, presence of food (left). Seeking without evaluating condition, absence of food (right). **c**, Representative single cell traces of LH^LepR^ neurons within several trials. **d**, 3-dimensional scored plot of LH^LepR^ neurons. **e**, Proportion of cell populations. **f**, **j**, Representative contour map of seeking (**f**) and evaluating (**j**) neurons. (top). Colour darkness represents cell activity degree (bottom). **g, k**, Representative single cell traces of LH^LepR^ neurons of seeking (**g**) and evaluating (**k**) neurons during food and no-food trials. **h, i, l, m,** Heatmap depicting the calcium signals (top) and average Z-scores (bottom) of seeking neurons (**h, i**) or evaluating neurons (**l, m**). Size of the calcium signals corresponds to its colour. (4 mice, 15 cells (**h,i**), 4 mice, 25 cells (**l,m**)). Data are mean ± s.e.m. See Supplementary Table 1 for statistics.

### Observed results of AgRP/NPY and LH^LepR^ neurons are consistent with the model of food need and motivation, respectively

Similar to our results on LH^LepR^ neurons, AgRP/NPY neurons are well known to evoke feeding behaviours when artificially stimulated and exhibit activity patterns related to each feeding bout^25^. To distinguish between the roles of LH^LepR^ and AgRP/NPY neurons, we compared theoretical modelling results based on the principles of homeostasis (Fig. 6a, b, Extended Data Fig. 6b, Supplementary Information) with observed neural activity data from experiments involving LH^LepR^ and AgRP/NPY neurons (Extended Data Fig. 6a). AgRP/NPY neurons were significantly inactivated at the food accessibility moment (Extended Data Fig. 6c–h) or when seeking had been initiated (Extended Data Fig. 6i–n)^31^. AgRP/NPY neurons were reactivated at the food inaccessibility moment (Extended Data Fig. 6o–t) or when mice terminated seeking or evaluating behaviour (Extended Data Fig. 6u–af). The observed AgRP/NPY neural activity was consistent with the theoretical (Fig. 6c) and computational (Fig. 6e) models of neural activity for “food need”, while the observed LH^LepR^ neural activity was consistent with the theoretical (Fig. 6d) and computational (Fig. 6i) models of neural activity for “food motivation”. Consistent with previous reports^20, 26^, mice displayed a latency to feeding when AgRP/NPY neurons were stimulated and exhibited significantly increased feeding behaviours that were sustained even after the termination of stimulation (Extended Data Fig. 7a-c, f-h, Supplementary Video 6). The behavioural patterns driven by the stimulation of AgRP/NPY neurons were in accordance with the computational model of behavioural activity for food need (Fig. 6g, Extended Data Fig. 7d, e, i, j). In contrast, the behaviours driven by stimulation of LH^LepR^ neurons exhibited a tightly time-locked temporal pattern without latency and abrupt cessation of feeding behaviour at the termination of stimulation (Supplementary Video 5). The behavioural patterns driven by the stimulation of LH^LepR^ neurons were in accordance with the computational model of behavioural activity for food motivation (Fig. 6k). Activation of AgRP/NPY neurons evoked all types of feeding behaviour, including seeking, evaluating, and swallowing, whereas activation of LH^LepR^ neurons significantly evoked only seeking and evaluating behaviours without a significant increase in swallowing (Supplementary Video 6). Collectively, our computational modelling results parsimoniously demonstrate that AgRP/NPY neurons encode food need, whereas LH^LepR^ neurons encode food motivation.

**Fig. 6.**
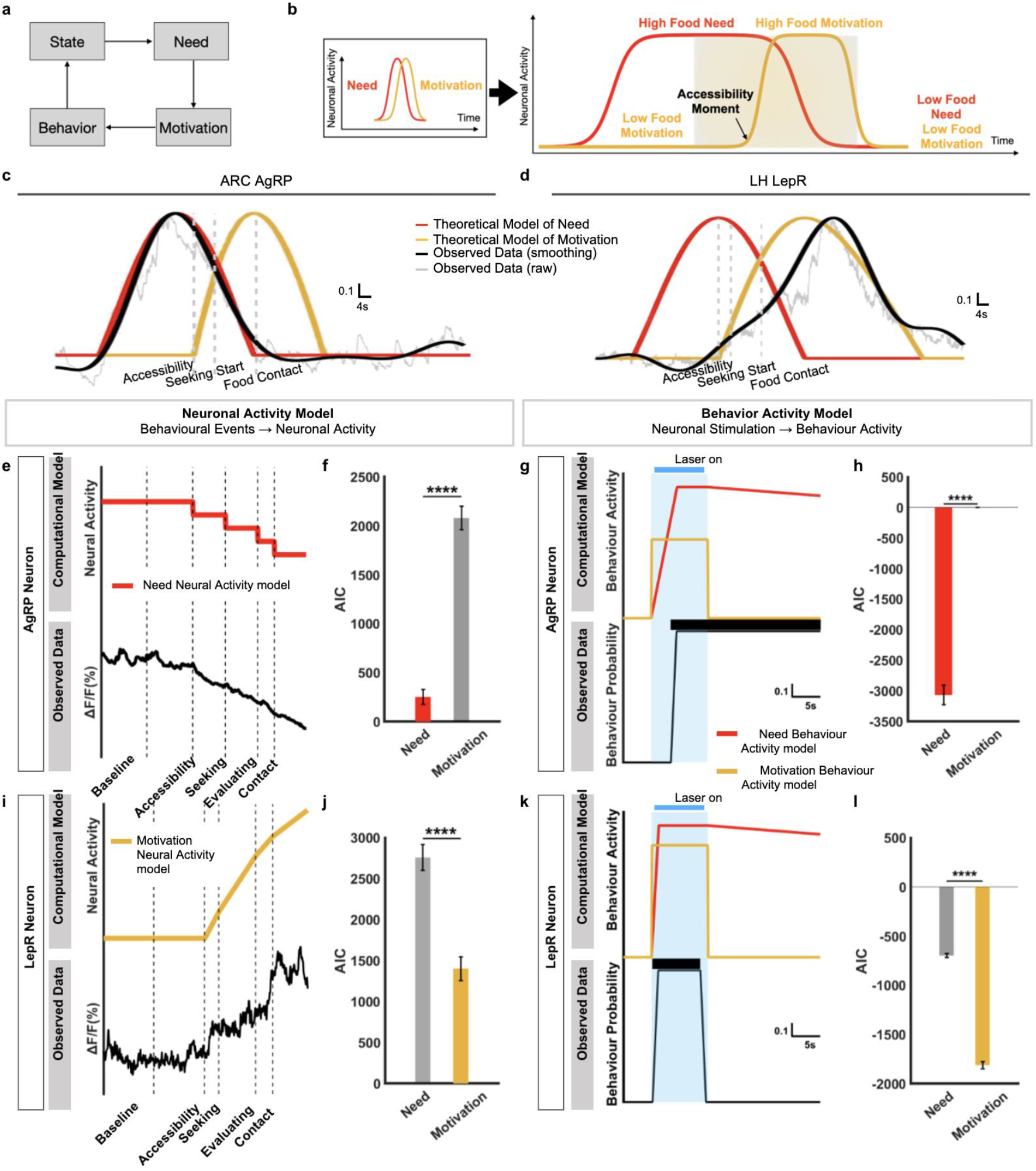
AgRP/NPY neurons and LH^LepR^ neurons are consistent with model of food need and motivation, respectively. **a**, Cycle of motivated behaviour. **b**, Activity of need (red) and motivation (yellow) in macro-timescale (left). Micro-scaled plot from left, distinguishable time window that dissect need and motivation (right). **c, d**, Theoretical model of need and motivation with observed AgRP/NPY (**c**) and LH^LepR^ (**d**) calcium signal. **e, f**, AgRP/NPY (**e**) and LH^LepR^ (**f**) neural activity comparison between computational model (left top) and observed neural activity (left down) with AIC score (right). **g, h**, AgRP/NPY (**g**) and LH^LepR^ (**h**) behavioural activity comparison between computational model (left top) and observed behaviour probability (left down) with AIC score (right).

### NPY increases LH^LepR^ neuron excitability via disinhibition of GABAergic interneuron in LH

To investigate how AgRP/NPY neurons encoding food need drive LH^LepR^ neurons encoding food motivation, we focused on NPY, which is the key modulator that generates sustained feeding behaviour after AgRP/NPY neural activity is inhibited ^25^. In addition, previous studies have reported that NPY receptors are distributed in the LH^27^ and that administration of NPY in the LH enhances feeding behaviour^28^.

To examine whether LH^LepR^ neurons respond to NPY, we performed slice calcium imaging in which artificial cerebrospinal fluid (ACSF) was applied to brain slices containing GCaMP6s-expressing LH^LepR^ neurons (Fig. 7a). LH^LepR^ neurons exhibited remarkably increased calcium activity after NPY application (Fig. 7b-d). The area under the curve (AUC) for the calcium signal and Z-score were significantly higher when NPY was applied than when ACSF was applied (Fig. 7e-f). Furthermore, the percentage of cells that responded to NPY was significantly higher than the percentage that responded to ACSF (Fig. 7g, h). Overall, these results suggest that NPY induces the excitability of LH^LepR^ neurons.

**Fig. 7.**
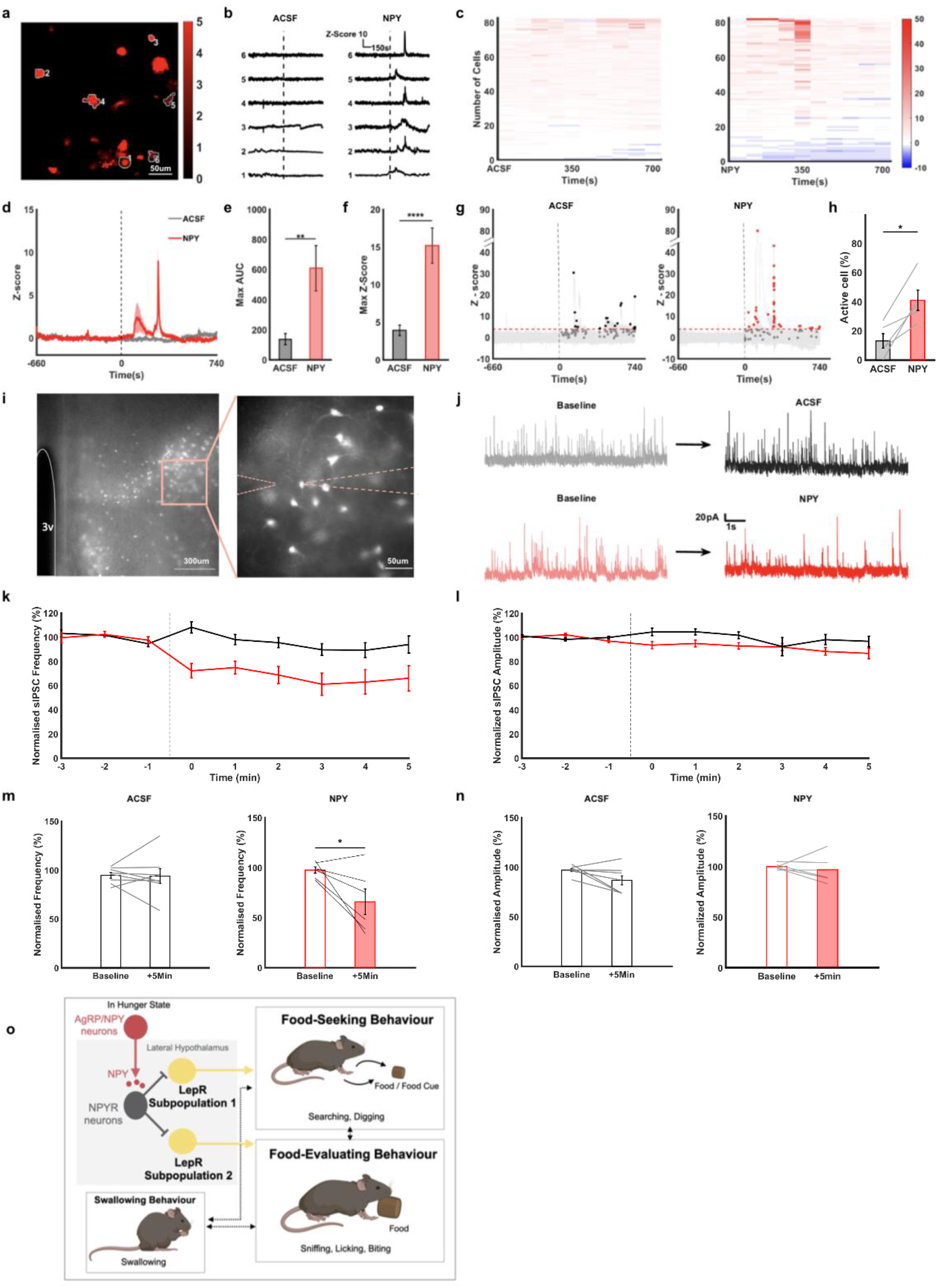
NPY increases LepR neuron excitability via disinhibition of GABAergic interneuron in LH. **a**, Representative image of GCaMP6s expressed LH^LepR^ neurons during brain slice calcium imaging. **b**, Representative traces of calcium activity of LH^LepR^ neurons marked in (**a**). Calcium activity of baseline and after bath application of ACSF or NPY. **c**, Absolute value of max Z-score of LH^LepR^ neurons in bins (each bin, 70s) after application of ACSF or NPY. **d**, Average traces of calcium activity after ACSF and NPY application. Dotted line is the time of application (4 slices each). **e, f**, Quantification of max AUC (**e**) and max Z-score (**f**) during (**d**). **g**, Individual traces of Z-score of calcium activity of LH^LepR^ neurons (grey traces). Black dot indicates max Z-score in responsive cells (17/94 cells) that activated (>4σ) after ACSF application. Red dot indicates max Z-score in responsive cells (48/94 cells) that activated (>4σ) after NPY application. **h**, Quantification of the percentage of active cells during (**d**). **i**, Representative images of td-Tomato-expressed LH^LepR^ neurons during brain slice whole-cell recording. **j**, Representative spontaneous inhibitory postsynaptic current traces comparing ACSF (top) and NPY (bottom). **k,l**, Time course of sIPSC frequency (**m**), amplitude (**n**) ( left) and quantification of normalised sIPSC frequency (**m**), amplitude (**n**) in the last 1min (right) after the ACSF or NPY application. (o) Comprehensive summary of neural circuits that encode need and motivation in feeding behaviour. Data are mean ± s.e.m. See Supplementary Table 1 for statistics.

Since NPY is reported to have an inhibitory effect on neural activity^29, 30^, we sought to determine the mechanism underlying the excitatory effect of NPY on LH^LepR^ neurons. According to our analysis of single-cell RNA sequencing data^22^ LH^LepR(-)/GABA(+)/NPYR(+)^ neurons accounted for a much larger proportion than LH^LepR(+)/GABA(+)/NPYR(+)^ neurons (Extended Data Fig. 1l). Therefore, we assumed that the excitatory effects of NPY in LH^LepR^ neurons arose from the disinhibition of presynaptic GABAergic neurons. To confirm whether the depolarisation of LH^LepR^ neurons resulted from a change in presynaptic input, we recorded inhibitory postsynaptic currents (IPSCs) in LH^LepR^ neurons before and after the application of NPY. (Fig. 7i, j). The frequency of IPSCs was significantly decreased by NPY application, but not with ACSF application (Fig. 7k, m). However, there were no changes in IPSC amplitude (Fig. 7l, n), suggesting an effect of NPY on presynaptic GABAergic neurons. Overall, these results suggest that NPY increases the excitability of LH^LepR^ neurons by decreasing inhibitory inputs.

## Discussion

The precise orchestration of need and motivation is critical for maintaining homeostasis^31^. Our findings demonstrated that LH^LepR^ neurons encode motivation for food-seeking and evaluating behaviours. AgRP/NPY neurons encoding the food need transmit information to LH ^LepR^ neurons using NPY as a permissive gate signal. (Fig. 7o).

To elucidate the exact role of LH^LepR^ neurons in feeding behavior, this study validated that the role of LH^LepR^ neurons involves intrinsic motivation. As a result, our data clearly show that the onset of LH^LepR^ neural activity precedes the onset of behavior, which suggests temporal causality of LH^LepR^ neurons as the motivation for the behavior. We show that LH^LepR^ neural modulation evokes food-motivated behavior through multiple food-seeking and evaluating tests. Also, we demonstrated that two distinct neuron populations exist among LH^LepR^ neurons, and each population separately encodes seeking and evaluating behavior to precisely orchestrate the appropriate feeding behavior.

The physiological need shows stepwise decrease when an animal perceives each predicted gain event (Supplementary Methods). In contrast, during these steps, motivation increases, and energy needs to be decreased in clearly opposite directions. Our model fitting analysis based on the theoretical homeostatic principle parsimoniously showed that AgRP/NPY neural activity is consistent with need-neurons and that LH^LepR^ neurons are consistent with motivation-neurons in feeding.

When the activity in AgRP/NPY neurons is high, NPY is sent to downstream regions to enhance feeding behaviour^20, 32^. Our *ex vivo* results indicated that NPY administration decreased the inhibitory input to LH^LepR^ neurons, suggesting that the excitability of LH^LepR^ neurons is increased and gated via decreases in tonic inhibition. This implies that LH^LepR^ neural activation is inhibited through tonic inhibition of upstream LH^LepR(-)/GABA(+)^ neurons during a low NPY state. In contrast, NPY inactivates LH^LepR(-)/GABA(+)^ neurons through NPY receptors during high NPY state. Of note, our *in vivo* results indicated that LH^LepR^ neuron activity began to increase after the presentation of food accessibility during a high NPY state.

Collectively, our *ex vivo* and *in vivo* results imply that NPY plays a permissive role in the activation of LH^LepR^ neurons. This putative role of NPY as an internal state-dependent permissive gate is adaptive for survival. In a satiated state, tonic inhibition functions as a lock, prohibiting LH^LepR^ neurons from responding to food-related cues. In the fasting state, NPY is released from AgRP/NPY neurons encoding hunger (food need), unlocking this tonic inhibition and allowing LH^LepR^ neurons to freely generate appropriate motivation in response to diverse food-related cues. This internal state-dependent conditional action of food motivation increases survival by restricting feeding behaviors only in the fasting state and by avoiding futile and unnecessary behaviors while the animal is sated.

The present study provides insight into how the brain encodes physiological needs and distinct motivations to orchestrate adaptive homeostatic behaviours. Examining the homeostatic mechanisms underlying food need and motivation will help to broaden our understanding of neuropsychiatric states such as obesity and addiction.

## Methods

### Animals

All experimental protocols were approved by Seoul National University Institutional Animal Care and Use Committee. Mice were housed on a 08:00 to 20:00 light cycle, with standard mouse chow and water provided ad libitum, unless otherwise noted. Behavioural tests were conducted during the light cycle. Adult male and female mice (at least 8-weeks-old) of the following strains were used: LepR-Cre (JAX stock no.008320), Vgat-IRES-Cre (JAX stock no.028862), Ai-14 Td-Tomato (JAX stock no.007914), and AgRP-IRES-Cre (JAX stock no.012899).

### Stereotaxic virus injection

Mice were anaesthetised with xylazine (20 mg/kg) and ketamine (120 mg/kg). A pulled-glass pipette was inserted into the arcuate nucleus (ARC) (300 nl total; AP, −1.3 mm; ML, ±0.2 mm; DV, 5.85 mm from the bregma) and the LH (400 nl total; AP, −1.5 mm; ML, ±0.9 mm; DV, 5.25 mm from the bregma) based on the 2D LH^LepR^ distribution (Extended Data Fig.1a-k). The GCaMP6 virus (AAV1.Syn.Flex.GCaMP6s.WPRE.SV40, Addgene; titre: 1.45×10^13^ genome copies per ml with 1:2 dilution) was utilised for calcium imaging. The AAV5.EF1⍺.DIO.hChR2(H134R).EYFP (Addgene; titre: 2.4×10^13^ genome copies per ml) or AAV5.EF1⍺.DIO.eNpHR3.0.EYFP (Addgene; titre: 1.1×10^13^ genome copies per ml) was utilised for optogenetic experiments. AAV8.hSyn.DIO.hM3D(Gq).mCherry (Addgene; titer 4.8×10^12^ genome copies per ml) was used for chemogenetic experiments.

### Optical fibre/GRIN lens insertion

For fibre photometry experiments, a ferrule-capped optical cannula (400 µm core, NA 0.57, Doric Lenses, MF2.5, 400/430–0.57) was unilaterally placed 0–50 µm above the virus injection site and attached to the skull with Metabond cement (C&B Super Bond). For optogenetic activation of AgRP/NPY neurons, a unilateral optical fibre (200 µm core, NA 0.37, Doric Lenses, ZF1.25_FLT) was implanted 100 µ m above the ARC injection site. For optogenetic manipulation of LepR neurons, optic fibres were bilaterally implanted 100–200 µm above the LH injection site at a 10° angle from the vertical in the lateral-to-medial direction. For micro-endoscope imaging, a GRIN lens (500 µm core, 8.4 length, Inscopix #1050-004413) was inserted after 3 weeks of recovery following virus injection. Dexamethasone, ketoprofen, and cefazolin were administered for postoperative care.

### Calcium imaging during feeding behavioural tests

#### Animal condition

Prior to experimentation, all mice were habituated to the experimental cages, and fibre handling was conducted for at least 3 days. During the behavioural experiments, all mice were housed in individual cages for food restriction, and their food was restricted to maintain 80–90% of the initial body weight in the fed state.

#### Voluntary behavioural maze test

This test mimicked the environment of a mouse in a cave, running to seek food despite the risk of fear outside, such as a predatory threat. When the motivation for food was higher than the fear of an electric shock, the mice made the voluntary decision to initiate seeking outside the doorless open shelter (Fig. 1j-k food test first column,1q, Extended Data Fig. 6i). On day 1, food zone conditioning was performed to allow mice to learn to eat the chocolate-flavoured snack at the end of the corridor until they were satiated. For the rest of the training sessions, trials were conducted with shock conditioning (random shock at a mean of 0.2 mA/shock with a 10-s interval). We adjusted the total duration of shock delivery to maximise the performance of mice.

#### Vertically placed food test

To measure total feeding behaviour or food-evaluating behaviour alone, we adjusted the height of the food tray. During obtainable height (8 cm) sessions, the mice engaged in both evaluating and swallowing behaviour (Fig. 1j-k food test, second column). During unobtainable height (11 cm) sessions, the mice engaged in evaluating behaviour only (Extended Data Fig. 3f, Extended Data Fig. 6aa).

#### Food or non-food test

The food or non-food test (Fig. 1j-k food test, third column, non-food test; Extended Data Fig. 2f) was performed using an L-shaped chamber in which food (chocolate-flavoured snacks) and inedible objects (Lego brick) were placed.

#### Food-conditioned maze test

The “seeking onset” was defined as the moment when the mouse left the shelter and ran straight to the food zone. The “evaluating onset” was defined as the moment when the mouse reached the top of the bridge and perceived visual food information. “Food contact” was defined as the moment when the mouse physically contacted food (Fig. 1m). To examine whether food accessibility influenced each behaviour, mice were either chased or pushed by force into the shelter, and the door was closed by the experimenter after the mice had received a chocolate-flavoured snack in the L-shaped chamber (Extended Data Fig. 6 o).

#### Visual cue maze test

During training, mice received chocolate-flavoured snacks when the food cue was visible. During testing, mice initiated seeking after presentation of the food cue. The success rate ([S2/W], S1 = number of abandonments, S2 = number of consumptions after food cue, W = S1+S2) was recorded during training until it reached 80%. The duration and amplitude of shocks during training were optimised for each mouse to achieve the best success rate (Extended Data Fig. 3a, Extended Data Fig. 6u).

### Neural manipulation during feeding behavioural tests

#### Animal condition

Prior to experimentation, all mice were habituated to the experimental cages, and fibre handling was conducted for at least 3 days. The mice had ad libitum access to water and food.

#### Hidden food-seeking test

For the conditioning session, chocolate-flavoured snacks or raisins were hidden under the wooden bedding at each edge of the field. Twice a day for 3 consecutive days, the mice were allowed to seek the box for hidden food during the 10 min of the experiment (Fig. 2c).

#### Nearby food-evaluating test

For DeepLabCut behavioural analysis (Fig. 2j-n), mice were acclimated in a transparent chamber (10 cm × 10 cm × 15 cm) with a socket to hold a cheese-flavoured snack inside. The test was recorded from below (ELP-USB4KHDR01-KV100, no-distortion camera) and from the side (Microsoft LifeCam HD-3000, no-distortion camera). During the testing session, laser stimulation (sham or real) was administered for 10 seconds under the above-mentioned conditions when the head of the mouse directly faced the food in the bottom and side views. Bottom and side views of the recorded videos were used for DeepLabCut analysis (24 frames per second). We labelled the snout, mouth (upper jaw, oral commissure, lower jaw), hands, paws, tail base, and food. We trained the network with 960 frames (in the bottom view) or 600 frames (in the side view) using a cut-off of 0.9 p for a total of 500,000 times. For manual behavioural analysis (Fig. 2o-p, Extended Data Fig. 7f-j), the period of non-locomotive evaluating behaviour (licking) was manually annotated while the tongue was visible. Compared with non-locomotive evaluating, wallowing was observed when the mice showed intense biting action.

#### Multiple food-evaluating test

On the test day, chocolate-flavoured snacks were given in an amount sufficient to ensure that the mouse would not eat all the snacks prior to the end of the test. On the test day, laser stimulation was delivered for 20 min at 2-min intervals (Fig 2s). Completion was defined as completing the swallowing of the snack, whereas incompletion was defined as premature termination of evaluating behaviour that resulted in remnant snacks left behind.

#### Food (or non-food) seeking and evaluating test

Cube sugar (food) and cube clay (non-food) were diagonally placed in each corner of an open-field box (30 cm × 30 cm × 30 cm) (Extended Data Fig. 4a). Laser stimulation was counterbalanced and delivered over 2 days.

#### Distant food-seeking test

At each side of the open-field box, a chocolate-flavoured snack or empty tray was placed at a height of 8 cm (Extended Data Fig. 7a). Laser stimulation was administered for 5 min.

### Fibre photometry

We used a Doric Lenses fibre photometry system. In the experiment, 465 nm and 405 nm LED light sources (Doric LED driver) were delivered continuously through a rotary joint (Doric Lenses, FRJ_1X1_PT-400/430/LWMJ-0.57_1m) connected to the patch cord (Doric Lenses, MFP_400/430/1100-0.57_1m), and the GCaMP6 signal was collected back through the same fibre into the photodetector (Doric Lenses).

### Optogenetics

Laser stimulation (473-nm, Shanghai BL473T8-150FC) was delivered through an FC-FC fibre patch cord (Doric Lenses) connected to the rotary joint, following which the FC-ZF 1.25 fibre patch cord delivered stimulation to the cannula (Doric Lenses, 200/220/0.23). The laser intensity for activation was approximately 10 mW at the tip.

### Chemogenetics

On the test day, clozapine-n-oxide (Tocris, CNO, 1 mg/kg, i.p.) or vehicle (saline, i.p.) was administered 1 hour before testing.

### 3D clearing

Fixed tissue was incubated in reflective index matching solution (C Match, Cat.50-3011) at 37℃ for 2 days. Images were obtained using SPIM (LaVision Biotech, Bielefeld, Germany) and analysed using IMARIS 9.5 (Bitplane AG, Zürich, Switzerland).

### Calcium imaging of brain slices

Brain slices were obtained from anaesthetised mice at least 3 weeks after virus injection. LH slices were dissected to a thickness of 250 μm using a vibratome (Leica, VT1200S) in ice-cold standard artificial cerebrospinal fluid (ACSF) containing the following (in mM): 125 NaCl, 2.5 KCl, 1 MgCl_2_, 2 CaCl_2_, 1.25 NaH_2_PO_4_, 26 NaHCO_3_, and 10 glucose, bubbled with 95% O_2_ and 5% CO_2_. For recovery, the slices were incubated at 32°C for 15 min and then further incubated for 1 h at room temperature. The slices were then transferred to the recording chamber and perfused with ACSF at 32°C during imaging. Calcium measurements were performed using a CMOS camera (Photometrics, USA) attached to an upright microscope (BX50WI, Olympus) with a 40X or 10X water-immersion objective (NA 0.8 or 0.3, LUMPlanFL N or UMPlanFl; Olympus) at 10 frames per second. A broad white light source (CoolLED, pE-340 fura, USA) was passed through an excitation filter (450–480 nm) and collected through an emission filter (525/50 nm). Fluorescence images were acquired using VisiView software (Visitron Systems GmbH, Germany).

### Single-cell RNA-seq analysis

scRNA-seq data with the LH (GSE125065) were analysed ^22^. Of the initial 7,232 cells (3,439 male and 3,793 female), 598 cells with less than 500 UMIs or >40% of mitochondrial reads were discarded. The R package Monocle3 was used to classify the cells^54^. Using Monocle 3, we subjected single-cell gene expression profiles to UMAP visualisation. Altogether, we identified 4,091 cells as neural clusters on the basis of cell type–specific marker gene expression ^22, 33, 34^. The neural clusters containing 4,091 cells were extracted for further clustering using Monocle 3 as above, which yielded 37 clusters. Clusters were classified as GABAergic if the median expression of Slc32a1 was greater than that of Slc17a6 in each cluster and glutamatergic if the median expression of Slc17a6 was greater than that of Slc32a1.

### Whole-cell patch clamp recording

Brain slices were obtained from aneasthetised (LepR-tdTomato) mice. LH slices were dissected to a thickness of 250 μm using a vibratome (Leica, VT1200S) with carbogen-saturated (95% O_2_ and 5% CO_2_) sucrose solution containing the following (in mM): 75 NaCl, 75 sucrose, 25 glucose, 26 NaHCO_3_, 7 MgCl_2_, 2.5 KCl, 1.25 NaH_2_PO_4_, and 0.5 CaCl_2_. For recovery, slices were incubated at 32°C for 15 min in standard ACSF containing the following (in mM): 125 NaCl, 2.5 KCl, 1 MgCl_2_, 2 CaCl_2_, 1.25 NaH_2_PO_4_, 26 NaHCO_3_, and 10 glucose, bubbled with 95% O_2_ and 5% CO_2_. Following further incubation for 1 h at room temperature, the slices were transferred to the recording chamber and perfused with ACSF at 32°C during recording. Whole-cell patch-clamp recordings were performed in LH neurons expressing tdTomato using EPC9 (HEKA). The resistance of the pipette was 2–5 MΩ when filled with an intracellular solution containing the following (in mM): 135 CsMS, 10 CsCl, 10 HEPES, 0.2 EGTA, 4 Na_2_-ATP, and 0.4 Na_3_-GTP (pH 7.2–7.3). All electrophysiological recordings were started at least 4 mins after the whole-cell configuration had been established.

### Drugs

Neuropeptide Y (NPY; Tocris) was dissolved in ACSF (400 µM) for slice application. The final drug concentration used in the calcium imaging experiments with bath application was 1 µM and that used in the whole-cell recording with pico-pump was 2 µM.

### Histology, immunohistochemistry, and imaging

Transcranial perfusion was performed using phosphate-buffered saline, followed by 4% neutral-buffered paraformaldehyde (T&I, BPP-9004). The brains were extracted, post-fixed in 4% paraformaldehyde at 4°C, and transferred to 10% sucrose, followed by 30% sucrose for cryoprotection. Cryoprotected brains were sectioned coronally on a cryostat (Leica Biosystems, CM3050) at 50 µm, and their sections were stained with 4’,6-diamidino-2-phenylindole (DAPI) to visualise the nuclei. To verify scientific exactitude, images of viral fluorescence and fibre/cannula placement were captured using a confocal microscope (Olympus, FV3000).

### Analysis

#### Behavioural tests

All data analyses were performed using custom-written MATLAB and Python codes. Behavioural experiments were manually analysed using Observer XT 13 or computationally analysed using EthoVision 14.

#### Quantification of neural onset

Neural onset can be determined by differentiating the neural activity recorded from calcium signals^35,36,37^. Neural activity was fitted to the optimal polynomial degree, and the third derivative of the fitted neural activity was calculated. Afterwards, the optimal time of the max value of the third derivative around the neural onset was determined.

#### Computational extraction of food-evaluating behaviour

To distinguish food-evaluating and swallowing behaviour from other behaviours, we first defined the duration in the food zone based on three criteria (Supplementary Video 4). In the bottom view, mice evaluating the cheese-flavoured snack stood up slightly, and the distance between the front paws decreased because they had been brought together to catch the cheese-flavoured snack. Therefore, the first criterion was met when the distance between the left and right front paws was less than that between the left and right hind paws in the bottom view. The second criterion was met when the y-coordinates decreased sequentially for the tail base, midpoint of the two hind paws, and midpoint of the two front paws in the bottom view. The third criterion was met when the snout coordinates were in the food zone in both the bottom and side views. To distinguish food-evaluating and food-swallowing behaviour, we defined food-swallowing based on two criteria: (1) presence of both front paws inside the food zone in the bottom view and (2) snout facing toward the centre of the food. The centre of the food was manually labelled in a randomly chosen frame from the test video and computed to change according to the distance ratio between the top, bottom, and centre of the food. Afterwards, the degree between two vectors was computed based on the snout and the centre of the food. The first vector was set from the snout to the centre of the food at a particular frame (N frame). The second vector was set from the location of the snout in the N and N+1 frames. The degree between two vectors was then smoothed using a moving average window of 1 s (24 frames). Finally, the second criterion was determined when the computed degree was between −20° and 20°. Food-evaluating behaviour was defined based on frames in which food swallowing behaviour did not occur.

#### Micro-endoscopic imaging

All data from the micro-endoscope experiments were recorded using nVoke (Inscopix). The raw signal output from CNMF-E (Craw) was converted into Z-scores (Z= (Craw-m)/σ), according to the mean (m) and standard deviation (σ) of the baseline (−10 s to −5 s before behavioural initiation). Activated cells were defined as cells with Z-scores of >4. Neural activity was normalised as follows: (NF0) = (Craw − minimum Craw) / (max Craw − minimum Craw).

#### Whole-cell Patch clamp recording

Spontaneous IPSCs were analysed using the Minhee Analysis Package^38^. The effects of NPY on sIPSCs were analysed by measuring the percentage change, compared to baseline for each neuron. The neurons in which the change was over 20% from the baseline were discarded, and those that exhibited sIPSCs over 5 Hz were used.

### Statistical analysis

All statistical data were analysed using MATLAB or IBM SPSS 25.0. Data in the figures are reported as the mean ± SEM. Paired t-tests were used to compare data between two groups. Two-way repeated-measures analyses of variance (ANOVA) were used for multiple comparisons. P-values for comparisons across multiple groups were corrected using the Greenhouse–Geisser method in IBM SPSS 25.0. Levels of significance were as follows: ∗p < 0.05. ∗∗p < 0.01, ∗∗∗p < 0.001, ∗∗∗∗p < 0.0001.

## Acknowledgments

We thank J.J.K. and M.J.Y. for analysing the calcium imaging data. This research was supported by the National Research Foundation of Korea (NRF) (NRF-2018R1A5A2025964, 2020R1C1C1012399 and 2021R1A6A3A13042965). Figures were generated using BioRender.

## Author Contributions

Y.H.L., Y.B.K., K.S.K., H.Y.S., M.J., and H.J.C. designed the project and interpreted the data. Y.H.L., Y.B.K., K.S.K., and M.J. wrote the paper with input from all authors. Y.B.K., Y.H.L., and H.Y.S. performed stereotaxic surgeries. Y.H.L., Y.B.K., H.Y.S., K.S.K., and D.-S.H. designed the behavioural tests. Y.H.L, Y.B.K., K.S.K., and H.Y.S. performed behavioural tests and data collection with contributions from D.-S.H., J.S.P., S.-H.J., K.M.K., and I.B. D.-S.H. and J.S.P. helped conceptualise the project. K.S.K. and Y.B.K. performed data analysis and visualisation with input from all authors. M.J. and J.L. performed brain slice experiments. D.C. performed 3D imaging. E.J.L. and M.S. analysed RNA sequencing data. S.J.K. and H.J.C. supervised the work. Y.H.L., Y.B.K., K.S.K., H.Y.S., and M.J. contributed equally to this work.

## Additional Information

Supplementary Information is available for this paper.

♦ Peer review information

## Data and code availability

The data and custom code that support the findings from this study are available from the corresponding author upon request.

## Supplementary Methods

### Theoretical computational modelling

#### Definitions

Homeostasis refers to the physiological process in which a stable state is maintained by a compensatory mechanism via an adaptive behaviour cycle^39, 40^. Energy state prediction is calculated by the expected loss or gain given that proactive behaviour is crucial for survival^41^. Predicted deficiencies are integrated into the motivation to evoke a particular behaviour^42, 43^. During the adaptive behaviour cycle, the predicted deficiency clearly decreases when an animal perceives the chance to earn a target (accessibility moment). The motivation to conduct a particular behaviour increases at this moment. Therefore, the activity of neurons encoding the predicted deficiency decreases while the activity of those encoding motivation increases in clearly opposite directions at this moment.

Need, which represents a predicted deficiency, fluctuates in response to a predicted gain or loss^44,45,46^. For example, the need for food will increase when travelling on an endless freeway with nothing to eat. However, the sight of a McDonald’s sign (accessibility moment) reduces the need even before entering the drive-through. Indeed, need arises from predicted deficiency, which is calculated by the sum of current deficiency and predicted gain or loss^40^ and starts to decline as the predicted gain begins to accumulate when accessibility is provided (seeing the McDonald’s sign). Need calculations based on predicted deficiency (rather than current deficiency) are more efficient for survival because calculations based on current deficiency induce over-compensation.

Motivation is derived from need, which is the force required to conduct a particular goal-directed behaviour^25, 42, 43, 47^. However, even when the need is high, motivation only starts to arise when the following prerequisites are met : absence of conflicting needs, no state of avolition, and (most importantly) accessibility to the goal is granted ^48, 49^. For example, during the freeway trip, the specific urge toward the hamburger increases only after observing the McDonald’s sign (accessibility moment). In addition, if the weather is too hot or the driver too thirsty, the driver will not feel the urge for food, despite energy deficiency and hunger.

Based on the principle of homeostasis, the current deficiency ***CD***(***t***_***r***_) is the represented as the deviation from the homeostatic state at a certain moment ***t***_***r***_ . This predicted deficiency ***PD*** (***t***_***r***_) is the sum of the current deficiency **CD**(***t***_***r***_) and the amount of predicted gain or loss induced by events ***E***(***t***_***r***_), which alleviates the state deviation z^39, 69^, as follows:

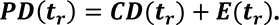

Event moments can be sequentially defined as (***t*_1_**, ***t*_2_**…***t***_***r***_ …***t*** _***n***_), where ***t***_***n***_ is the moment when an animal reaches its goal. The sum of the predicted gain or loss is based on past events perceived in the context^70^. As events occur in series, the predicted gain or loss accumulates until the goal is attained^50^. Therefore, the predicted deficiency at ***t***_***r***_ can be redefined as the sum of the current deficiency and accumulation of all predicted gains or losses:

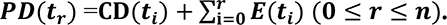

Need is the valuation of the predicted deficiency ***V***(***PD*** ((***t***_***r***_)).^41, 51^ Therefore, the need at ***t***_***r***_ can be expressed as follows:

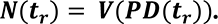

Motivation is the integration of needs through accumulation^3, 44, 61, 64^. Motivation is affected by an integrated affecting factor (**a**). For example, motivation will not accumulate before accessibility is granted^49^ (**a** = **0**). Motivation will also change according to factors such as satiety, valence, and memory^5, 12, 49, 52, 53^. Therefore, motivation at a certain time point (**M**(***t***_***r***_)) can be written as the product of the integrated affecting factor and the integral of the following:

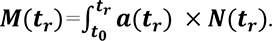

When sufficient motivation is accumulated to pass a certain threshold (K ≥ 0), behaviour begins to appear. Behaviour is reinforced according to the accumulation of motivation. Therefore, behavioural activity at a certain time point (***t***_***r***_) can be expressed as follows:

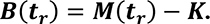

### Neural activity model

To decode the optimal model for each experiment, each trial from all experiments was extracted according to its initial behaviour annotation time point. The need value in each experiment was defined as the difference between the maximum and minimum z-scores across the extracted time period.

Based on this formula, the need was computed to sharply decrease at the behaviour moments annotated in each test. The motivation was computed as 0 before the annotated accessibility moment.

Afterwards, motivation was computed as the integration of need, and the integrated affecting factor was defined as the reciprocal of the maximum z-scores for optimal fitting. Inhibition and activation models were designed to strictly decrease or increase at a specific behavioural event to the maximum or minimum z-score value for the given test and to be sustained until trial termination or the occurrence of a plausible change.

Experimental data were fitted to fifth-order polynomials for comparison with the computed model. All computational data derived from the experimental data were aligned to the mean of the baseline period of the experimental data for equivalent comparison across the derivate models. To find the optimal model for each experiment and trial, the Akaike information criterion (AIC) was used to quantify the goodness of fit of the models when compared with the experimental data^76^. The AIC values were calculated as follows:

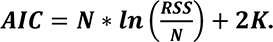

N is the number of data points, RSS is the residual sum of squares, and K is the number of free parameters. Differences between models were compared by calculating the differential AIC between the base models and other models (ΔAIC = AIC_other_ − AIC_base_). Recently, the computation of AIC for in *vivo* calcium signals in dopamine neurons demonstrated a unified interpretation of their role in learning^54^. Paired t-tests were performed to compare ΔAIC values.

### Behavioural activity model

To decode the optimal model for each experiment, the probability of behavioural activity was calculated and then decoded in a pipeline.

The probability of behavioural activity was computed from periodic annotations of mouse behaviour using a timespan of 1 s split into bins (100 bins for distant food-seeking test) (Extended Data Fig. 7d-e) and 10 bins for the nearby food-evaluating test (Extended Data Fig. 7i-j). The bin was recorded as “1” if a behaviour occurred during the period for that bin. Afterwards, all time-split trials in the test were averaged and normalised to determine the probability of behavioural activity. The behavioural activity for each time bin was then calculated with a moving average (window of 10 bins for tests with 10 s of stimulation, 60 bins for the rest) to obtain a smooth curve for comparison.

To identify the optimal model for behavioural activity in each experiment, we generated models for neurons encoding need and motivation. In these models, the need value in the behavioural activity model of need neurons was defined as 1 during neuronal activation. At the stimulation offset, the need value was computed to decrease strictly to an estimated constant degree (1/300 per second) for optimal decay modelling of the accumulated motivation level. The motivation level was computed as accumulation of the computed need and normalised proportionally according to the maximum value of the experimental data. After normalisation, motivation was computed to be sustained until stimulation offset when the motivation level reached the first highest peak of observed behaviour activity. Behaviour activity was then computed to increase according to the motivation level when the motivation level passed a certain threshold (K>0).

In the model used to determine the probability of behavioural activity, the motivation value was computed to be sustained at 0 until the onset of neuronal activation. During neuronal activation, the onset motivation value was computed to rise sharply. Behavioural activity was then computed to perpendicularly rise to a certain value after motivation had passed a certain threshold and was sustained horizontally until the offset of stimulation. Behavioural activity was then computed to increase according to the motivation level when the motivation level had passed a certain threshold (K>0). To determine the minimum AIC, the AIC was calculated while changing the motivation value in increments of 0.05. The values from all models were calculated using the ΔAIC, and analysis was performed as stated above for further comparison.

## Supplementary Video Legends

Supplementary Video 1. LHᴸᵉᵖᴿ 3D mapping, related to Extended Data Figure 1.

Supplementary Video 2. LH^LepR^ neuron activity is time-locked to food-seeking and evaluating.

The grey line indicates LHᴸᵉᵖᴿ neural activity during behavioural tests. The black line indicates neural activity fitted to the 9th order polynomial. The yellow line indicates the maximum value of the third derivative of neural activity (jolt) used to calculate neural onset. 00:00-03:17 shows that LHᴸᵉᵖᴿ neural activity increases during food-seeking and evaluating. 00:04-01:03 shows the vertical food placement test (obtainable condition). 01:04-03:03 shows the food-conditioned maze test. 03:04-03:17 shows the voluntary behaviour maze test. 03:18-03:40 shows that LHᴸᵉᵖᴿ neural activity starts decreasing at the beginning of food-seeking and termination of evaluating behaviour (vertically food placed test, unobtainable).

Supplementary Video 3. LH^LepR^ neurons drive motivation for food-seeking and evaluating behaviours. 00:00-01:32 shows that activation of LHᴸᵉᵖᴿ neurons drives motivation for food-seeking and evaluating. 00:03-01:12 shows the hidden food-seeking test. The yellow box indicates the seeking zone. 01:13-01:32 shows the nearby food-evaluating test. 01:33-03:35 shows that inhibition of LHᴸᵉᵖᴿ neurons decreases motivation for food-evaluating behaviour (food-evaluating test). The black box indicates the duration of evaluating behaviour. If the mouse completed evaluating behaviour, a blue line was displayed at the end of the black box. If the mouse did not complete evaluating, a red line was displayed at the end of the black box.

Supplementary Video 4. Classification of food-evaluating and swallowing behaviours using DeepLabCut. The video above shows the degree between two vectors, which was used to define the second criterion (Methods). The blue line with the arrow indicates normalisation for optimal visualisation. The video below shows an analysis of coordinates fulfilling the first criterion (Methods). 00:04-00:16 shows food-evaluating behaviour. 00:17-00:49 shows food-swallowing behaviour.

Supplementary Video 5. Representative calcium activity of LHᴸᵉᵖᴿ neurons during food-seeking and evaluating behaviour.

The upper right area shows activated LHᴸᵉᵖᴿ neurons. Darker colour represents higher cell activity. 00:03-00:25 shows a food trial (seeking with evaluating). 00:26-00:50 shows a no-food trial (seeking without evaluating).

Supplementary Video 6. Comparison of neural activity in AgRP/NPY neurons and LHᴸᵉᵖᴿ neurons and the associated behavioural patterns.

00:00-00:52 shows that activation of AgRP/NPY neurons evokes all types of feeding behaviour, whereas activation of LH^LepR^ neurons evokes only evaluating behaviour. 00:53-01:22 shows that activity in AgRP/NPY neurons and LHᴸᵉᵖᴿ neurons is consistent with the theoretical models of food need and motivation, respectively. The red line represents neural activity for the theoretical model of need neural activity. The yellow line is theoretical model of motivation neural activity. The grey line is neural activity measured via fibre photometry. The black line indicates the neural activity fitted to a 20^th^-degree polynomial.

---

**Extended Data Fig. 1.**
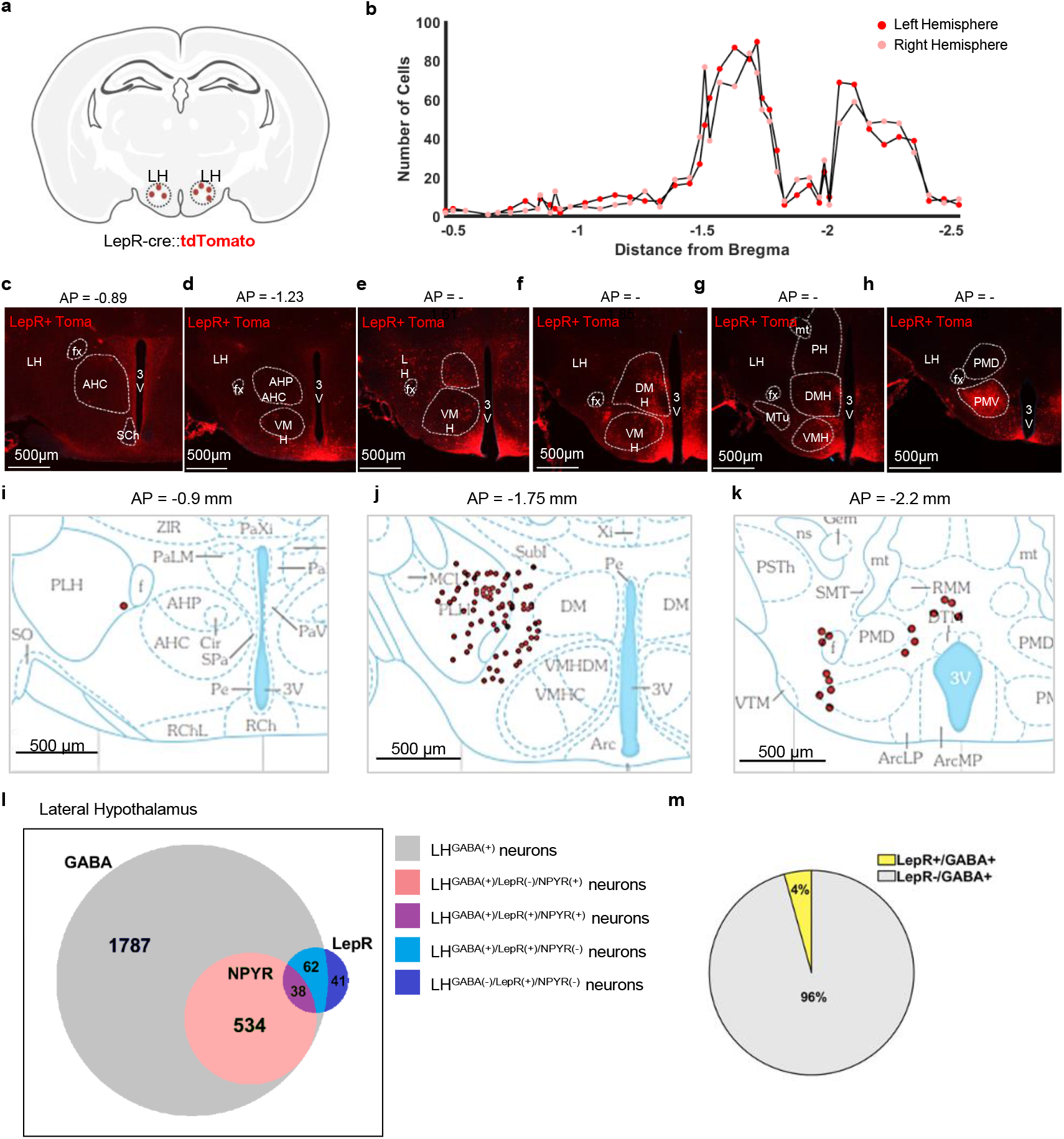
Distribution and molecular identity of LepR neural population in the lateral hypothalamus. **a, b**, Distribution profile and quantification of LepR-positive cells in the LH along the anterior-posterior axis of LepR-tdTomato mice (3 mice). **c-h**, Coronal brain sections showing tdTomato+ cells. **i-k**, Representative image depicting the distribution of LH^LepR^ tdTomato+ cell bodies. **l**, Venn diagram of molecular characteristics (GABA, NPYR, LepR) of the LH neurons by single-cell RNA sequencing data. **m**, Proportion of LepR-positive (yellow) and LepR-negative (grey) among GABA-positive neurons by single-cell RNA sequencing data.

**Extended Data Fig. 2.**
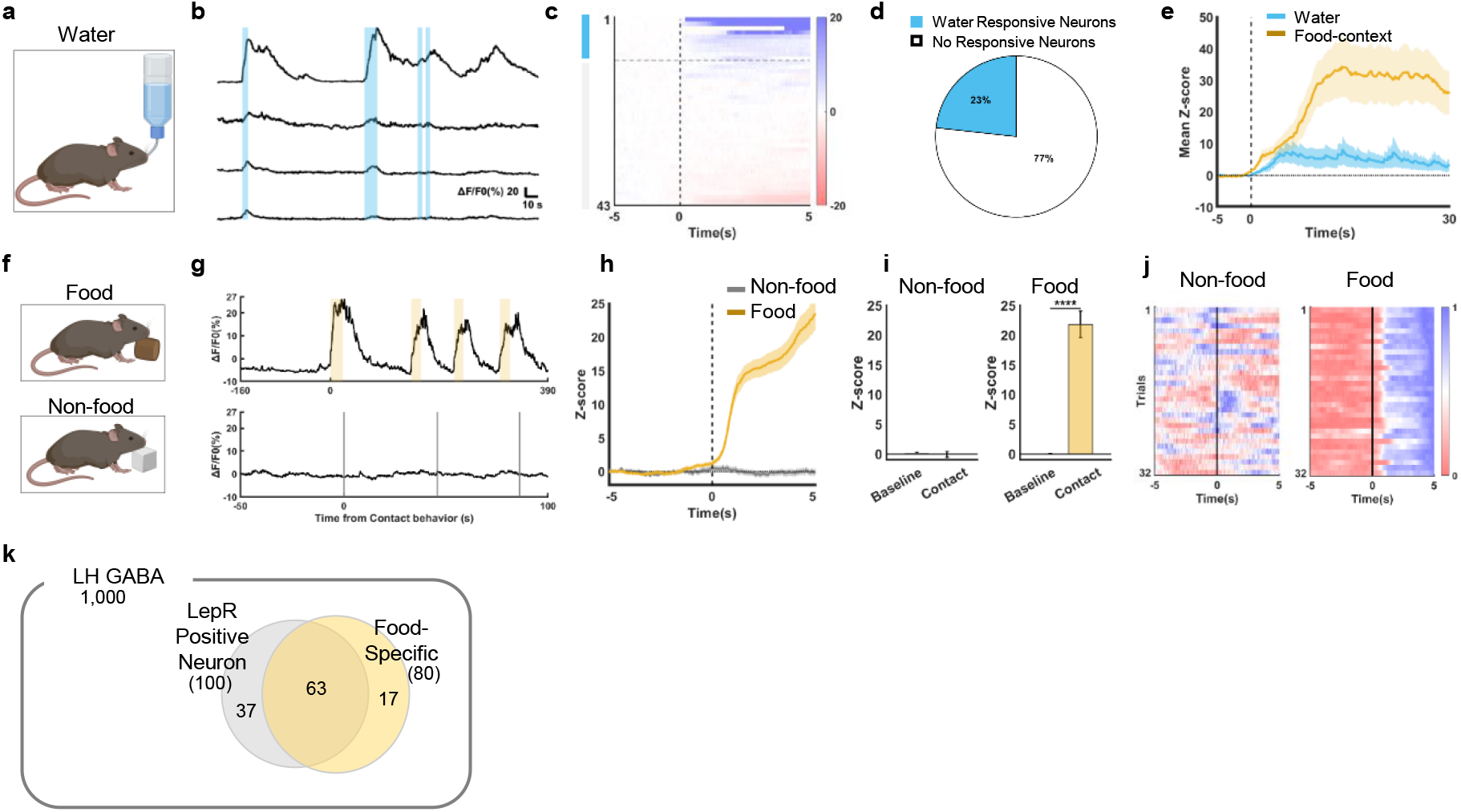
Activity of LH^LepR^ neurons is food-specific. **a**, Schematic of the behavioural test for drinking water in the dehydration state. **b**, Representative single-cell traces of LH^LepR^ neurons aligned with drinking behaviour. Blue shaded box indicates each bout of drinking. **c**, Heatmap depicting calcium signals that respond to water. Water-responsive cells (blue) activated (>4σ) during drinking water. **d**, Proportion of water-responsive cells (blue, 23%) and no responsive cells (white, 77%). **e**, Average Z-score from LH^LepR^ calcium signal aligned to drinking behaviour for water (blue) and feeding behaviour for food (yellow). **f**, Schematic of food/non food test. **g**,**h**, Representative calcium traces (**g)** and average Z-score (**h)** from the LH^LepR^ calcium signal aligned to contact of food (top) and non-food (below). The yellow shaded box indicates behaviour from food contact to end of consumption, and grey line indicates contact to non-food. **i**, Quantification of the Z-score in calcium signal changes from (**h**). Comparison between the baseline (−8 to −7s) and after contact (4 to 5s). (4 mice; 32 trials). **j**, Heatmap depicting normalised LH^LepR^ calcium signal aligned to contact with food and non-food. **k**, Venn diagram simulating the number of LH^LepR^ -positive and food-specific neurons when the total number of LH ^GABA^ neurons is simulated as 1,000. Data are mean ± s.e.m. See Supplementary Table 1 for statistics

**Extended Data Fig. 3.**
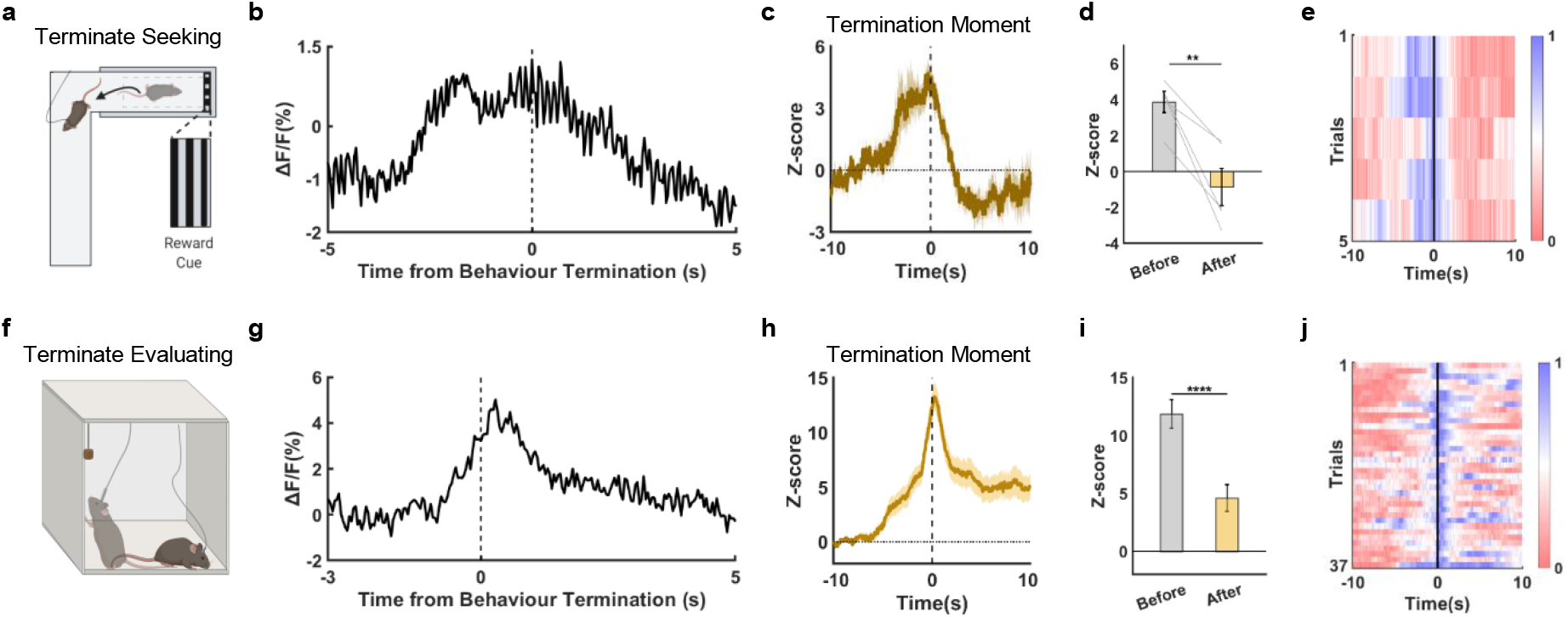
LH^LepR^ neurons are inactivated at the termination moment of food-seeking and evaluating. **a**, **f**, Schematic of the visual cue maze test (**a**) and vertically placed food test (unobtainable) (**f**). **b**, **g**, Representative calcium signal of LH^LepR^ neurons aligned to the termination moment of seeking (**b**) and evaluating (**g**). **c**, **h**, Average Z-score from LH^LepR^ neurons aligned to the termination moment of seeking (**c**) and evaluating (**h**). **d**, **i**, Quantification of the Z-score from (**c**, **h**). Comparison between the before (0 to 1s) and after behavioural termination (9–10s). (**d**) (1 mouse, 5 trials) (**i**) (5 mice, 37 trials). **e**, **j**, Heatmap depicting normalised LH^LepR^ calcium signal aligned to the termination moment of seeking (**e**) and evaluating behaviour (**j**). Data are mean ± s.e.m. See Supplementary Table 1 for statistics.

**Extended Data Fig. 4.**
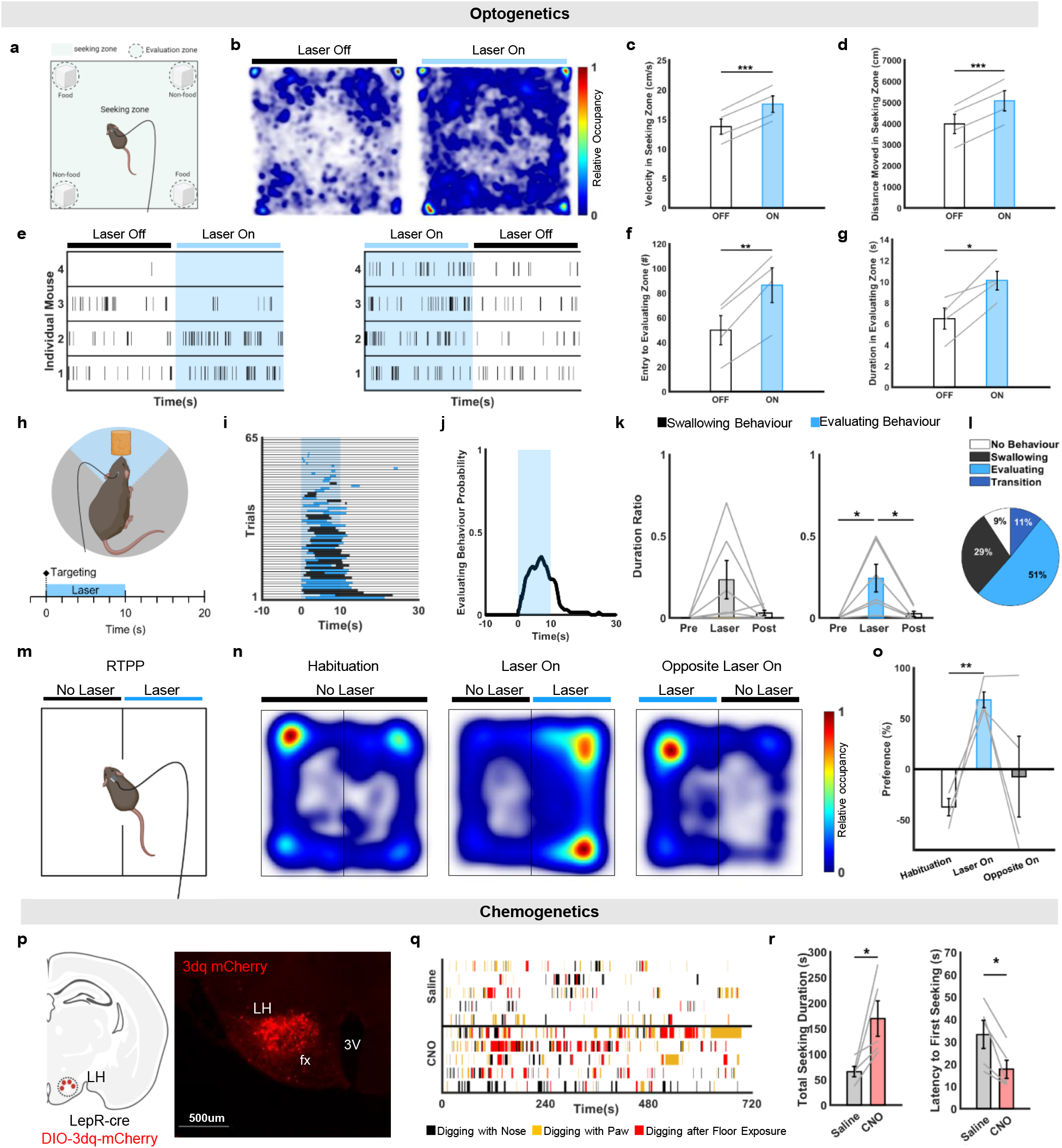
Activation of LH^LepR^ neurons drives food-seeking and evaluating behaviour. **a**, Schematic of the food/non-food-seeking and evaluating test. **b**, Representative occupancy heatmaps of time spent during (**a**). **c**, **d, f, g**, Quantification of velocity (**c**), distance moved (**d**) in the seeking zone, quantification of entry (**f**) and duration (**g**) of the evaluating zone (4 mice). **e**, Raster plot of food-seeking behaviour. Laser stimulation was delivered in a counterbalanced manner. (4 mice). **h**, Schematic of the nearby food-evaluating test. The laser is stimulated when the head of the mice is in the food zone (blue). **i**, **j**, Raster plot (**i**) and behavioural probability (**j**) of evaluating (6 mice). **k**, Quantification of swallowing behaviour (left) and evaluating behaviour (right) from (**i**) (6 mice). **l**, Proportion of feeding behaviours from (**i**) (6 mice). (**m**) Schematic of the real-time place preference test. Black bar indicates unpaired zone, and blue bar indicates photostimulation-paired zone. (**n**) Representative occupancy heatmaps of time spent in each zone during habituation (left), photostimulation (middle), and opposite-side photostimulation (right). (**o**) Quantification of preference for each zone during habituation (white), photostimulation (blue), and opposite-side photostimulation (gray). (4 mice) **p**, Schematic of chemogenetic activation and image of 3dq expression in the posterior LH^LepR^ neurons. **q**, Raster plot of seeking after injection of saline (top) and CNO (bottom). **o**, Quantification of seeking during (**q**) after injection of saline (grey) and CNO (pink). Total seeking duration(left), and latency to first seeking (right). (n=5). Data are mean ± s.e.m. See Supplementary Table 1 for statistics.

**Extended Data Fig. 5.**
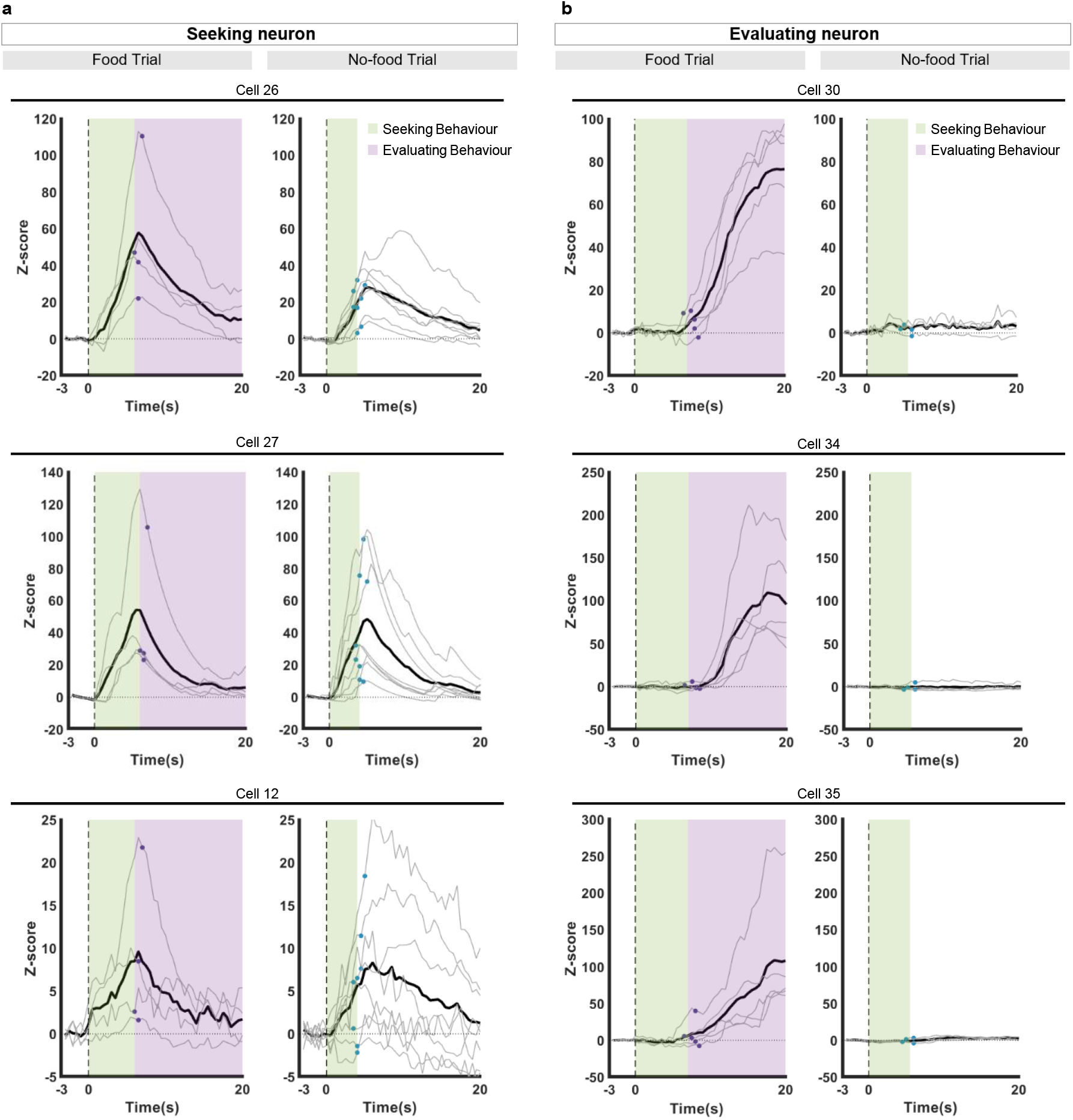
Seeking and evaluating neurons are separated in individual traces of LH^LepR^ neurons. **a**, **b**, Representative activity of an individual neuron from seeking-neurons (**a**) and evaluating-neurons (**b**). Average Z-score (bold) aligned to the seeking initiation moment. Grey lines indicate calcium signal of individual trials. Violet dot is the moment of physical food contact (initiation of evaluating). Blue dot is the moment of entry into the food zone. Green shade is seeking phase and purple shade is evaluating phase. Data are mean ± s.e.m. See Supplementary Table 1 for statistics.

**Extended Data Fig. 6.**
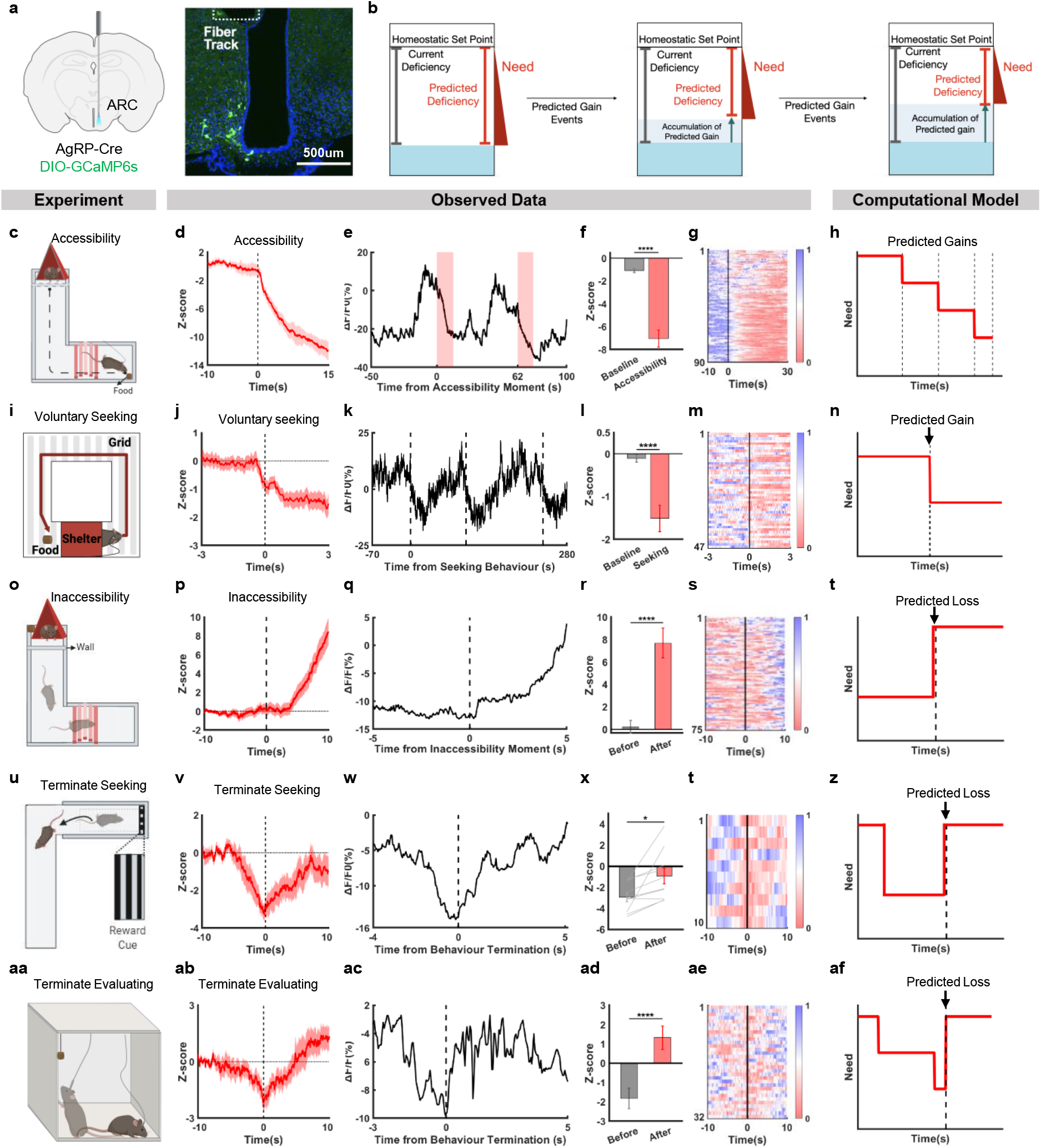
AgRP/NPY neural activity is consistent with need neural activity model. **a**, Schematic of virus injection/fibre insertion for fibre photometry in the arcuate nucleus from AgRP-Cre mice. **b**, Schematic of change in physiological need according to the process of decreasing the predicted deficiency and the accumulation of predicted gain. **c**, **i**, Schematic of the predicted gain tests with the accessibility (**c**) and voluntary seeking initiation moment (**i**). **d**, **j**, Average Z-score from the AgRP/NPY neural activity response to predicted gain (aligned to accessibility (**d**) or voluntary seeking initiation moment (**j**)). **e**, **k**, Representative AgRP/NPY neural activity traces response to predicted gain (aligned to accessibility m (**e**) or voluntary seeking initiation moment (**k**)). **f**, **l**, Quantification of Z-score in calcium signal changes from (**d**, **j**). Comparison between the baseline (−8 to −7s) and after accessibility (14 to 15s) (**f**) and after voluntary seeking (2 to 3s) (**l**). **g**, **m**, Heatmap depicting normalised AgRP/NPY neural activity aligned to accessibility (**g**) or voluntary seeking initiation moment (**m**). **h**, **n**, Computational need neural activity model (red bold line) based on predicted events. The dashed lines indicate the moment of predicted events initiation. **o**, **u**, **aa**, Schematic of the predicted loss tests with the inaccessibility (**o**), termination of seeking (**u**), and termination of evaluating moment (**aa**). **p**, **v**, **ab**, Average Z-score from AgRP/NPY neural activity response to predicted loss (aligned to inaccessibility (**p**), termination of seeking (**v**) or termination of evaluating (**ab**)). **q**, **w**, **ac**, Representative AgRP/NPY neural activity response to predicted loss (aligned to inaccessibility (**q**), termination of seeking (**w**) or termination of evaluating (**ac**)). **r**, **x**, **ad**, Quantification of Z-score in calcium signal changes from (**p**, **v**, **ab**). Comparison between the before (0 to 1s) and after (9 to 10s). (5 mice, 75 trials (**r**), 2 mice, 10 trials (**x**), 4 mice, 32 trials (**ad**). **s, y, ae,** Heatmap depicting normalised AgRP neural activity aligned (aligned to inaccessibility (**s**), termination of seeking (**y**) or termination of evaluating (**ae**)). **t**, **z**, **af**, Computational need neural activity model (red bold line) based on predicted gain/loss events, aligned to predicted loss (**t, af**). The dashed lines indicate the moment of predicted events initiation. Data are mean ± s.e.m. See Supplementary Table 1 for statistics.

**Extended Data Fig 7.**
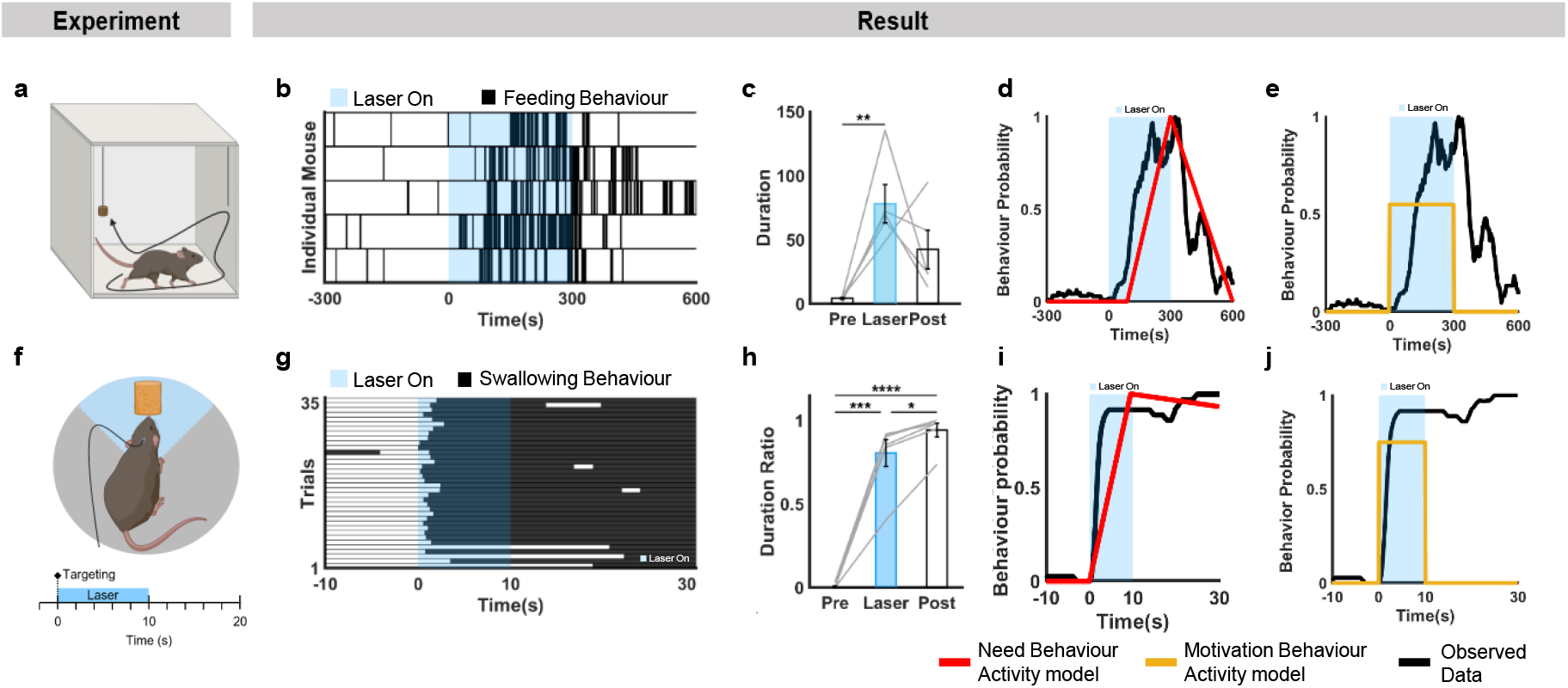
Behavioural activity of AgRP/NPY optogenetic stimulation is consistent with need behavioural activity model. **a**, Schematic of the distant food-seeking test. **b**, Raster plot of feeding behaviour during (**a)**. **c**, Quantification of the duration of feeding behaviour during pre-stimulation, stimulation, and post-stimulation (5 mice). **d, e**, Comparison between the observed data and the two computational behavioural activity models. The probability of behavioural activity (black bold line) and need behavioural activity model (red bold line) (**d**) and motivation behavioural activity model (yellow bold line) (**e**). **f**, Schematic of the near food-evaluating test. The laser is on when the head of the mice is in the food zone. **g**, Raster plot of feeding behaviour during (**f**). **h**, Quantification of the duration of feeding behaviour during pre-stimulation, stimulation, and post-stimulation (5 mice). **i**, **j**, Comparison between the observed data and the two computational behavioural activity models. The probability of behavioural activity (black bold line) and need behavioural activity model (red bold line) (**i**) and motivation behavioural activity model (yellow bold line) (**j**). Data are mean ± s.e.m. See Supplementary Table 1 for statistics.

